# Cfap410a and Cby work together with tissue-specific requirements to build *Drosophila* ciliary transition zones

**DOI:** 10.64898/2026.09.08.749861

**Authors:** Amélie Billon, Jennifer Vieillard, Pranjali Priya, Minita Desai, Emilie Fontaine, Sarah de Freitas, Jean-André Lapart, Alain Mangé, Haneef Ahmad Dar, Véronique Morel, Joëlle Thomas, Swadhin Chandra Jana, Bénédicte Durand

## Abstract

Cilia and flagella perform essential physiological functions in eukaryotes, and defects in these organelles cause several human diseases, including cancer and ciliopathies. The architecture of cilia is highly organized. The ciliary compartment is separated from the cytoplasm by the transition zone (TZ). The severity of ciliopathies linked to TZ assembly defects highlights the TZ’s critical role. Although several core conserved complexes are involved in TZ assembly, variations in TZ composition are associated with structurally and functionally diverse cilia. Here, we identify Cfap410a as a novel component of the ciliary TZ in the two *Drosophila* ciliated tissues, male germ cells and sensory neurons. Cfap410a is one of the two *Drosophila* paralogs (Cfap410a and Cfap410b) of human CFAP410, whose mutations are associated with axial spondylo-metaphyseal dysplasia, retinitis pigmentosa and amyotrophic lateral sclerosis. We show here that Cfap410a is a proximity partner of Cby and that they act cooperatively in the hierarchy of the TZ assembly program by bridging the CEP290 and MKS transition zone modules. Simultaneous loss of Cfap410a and Cby halts ciliary growth by disrupting TZ formation in multiple types of *Drosophila* ciliated cells, each of which exhibiting varying dependence on these two proteins. Interestingly, the function of Cfap410a and Cfap410b are not functionally redundant, indicating that the two proteins have evolved towards specific functions. In summary, our results propose a novel role for CFAP410a at the TZ and provide an explanation for how deregulation of conserved TZ components could lead to tissue-specific ciliopathies.

## Introduction

Cilia and flagella are highly conserved organelles among eukaryotes. They protrude at the surface of almost all mammalian cells and play numerous and essential physiological roles in our body. Any defects in their assembly or function, lead to various, and sometimes lethal diseases, collectively named ciliopathies.

Cilia are built around a conserved microtubule scaffold, the axoneme, surrounded by the ciliary membrane. At the ciliary base lies a highly organized structure, the transition zone (TZ), which connects the basal body (BB) to the plasma membrane and controls protein movement in and out of the cilium. The critical role of the TZ is revealed by the severity of the ciliopathies caused by TZ assembly defects [1,2].

Several protein complexes have been described to be involved in TZ assembly, among which the Cep290, MKS and the Joubert modules, conserved in most ciliated eukaryotes [3]. The hierarchical spatial organization of these complexes have been described and compared in several organisms [3–6]. Cep290 also recruits another set of proteins, the Dzip1-Cby-Fam92 complex which is critical for building the TZ in *Drosophila* and vertebrates [7,8]. But only Dzip1 is conserved from protozoa to mammals, whereas Cby and Fam92 are not found in *C. elegans* or protozoa [8,9], illustrating variations between TZ assembly modules in different organisms.

Defects in TZ components in humans lead to specific ciliopathies including the Joubert syndrome, Meckel-Gruber syndrome and retinopathies. In Human, *CFAP410* mutations have been associated with retinitis pigmentosa (RP) [10], rod cone degeneration [11,12], skeletal ciliopathies [13–15], the Joubert syndrome [13] and also with the neurodegenerative disease amyotrophic lateral sclerosis (ALS) [16–18]. The first four diseases are related to ciliary dysfunction but the mechanisms by which CFAP410 regulates cilium assembly or function is largely unknown, despite a precise characterization of the structure of its N and C-terminal domains in *Trypanosoma* [19,20]. In retinal cells, CFAP410 was shown to be associated with the periciliary compartment adjacent to the daughter centriole, but also with the connecting cilium of the photoreceptor [10,14], which is proposed to be a modified transition zone as it harbors features of the ciliary TZ. Since the Joubert syndrome is associated with defects of TZ components, this supports the hypothesis that CFAP410 plays a role at the TZ. However, the molecular mechanisms supporting CFAP410’s roles in the various diseases are still unknown.

The *Drosophila* genome contains two *CFAP410* orthologs: *cfap410a*/*CG14995* and *cfap410b*/*CG15208*. We identified Cfap410a as a novel component of the ciliary transition zone in the two types of *Drosophila* ciliated cells: male germ cells and sensory neurons. Cfap410a and Cfap410b do not compensate for each other, showing that these homologous proteins have distinct functions. *Drosophila* Cfap410a links the CEP290 and MKS modules of the transition zone. Cfap410a interacts with Cby to facilitate transition zone assembly. Losing both proteins disrupts transition zone integrity and prevents cilia formation with specific extent in the different *Drosophila* ciliated cell types.

## Results

### CFAP410 is conserved in ciliated organisms and *Drosophila* Cfap410a is a TZ protein in all ciliated tissues

CFAP410 orthologues are found in many ciliated organisms and various number of paralogs can be found. One ortholog is found in *C. elegans* (F09G8.5), *Trypanosoma* and *Chlamydomonas* [19,20] and up to 6 in *Paramecia* (Figure S1A and B). Alignment of proteins from *Paramecia*, *C. elegans*, *Drosophila*, mouse and Human shows a highly conserved region in the N-terminal domain, which exhibits four Leucine Rich Repeat domains of unknown function that adopt a typical helix conformation, as predicted by Alphafold [21,22] and the crystal structure of this domain is described in *Trypanosoma* [19]. There is also a small conserved helix domain at the C-terminal (Figure S1A). Recent structural studies showed that this C-terminal domain is involved in protein tetramerization and basal body localization in *Trypanosoma* [20]. *cfap410a/CG14995* and *cfap410b/CG15208* are the only two *Drosophila* orthologs of Human *CFAP410*.

To determine the localization of the two proteins in flies, we generated transgenic *Drosophila* lines that expressed each of the fusion proteins, Cfap410a-Tom or Cfap410b-GFP, under the promoter of their respective gene. We detected Cfap410a in the ciliary compartment of male germ cells and Type-I sensory neurons. In germ cells, Cfap410a first appeared in G2 spermatocytes. At this stage, there are two pairs of centrioles which convert into four basal bodies (BB) that dock to the plasma membrane, where they form an elongated TZ or primary cilium like (Figure 1A). During meiosis, the BBs stay attached to the plasma membrane that invaginates inside the cytoplasm, thus maintaining a ciliary cap/TZ at their distal ends. After meiosis, the BB anchors to the nucleus via its proximal end, the axoneme starts assembling inside the ciliary cap, while the TZ/ring centriole detaches from the BB and migrates along the elongating axoneme, keeping the integrity of the ciliary cap (Figure 1A) [23–27]. We show that Cfap410a appears at centrioles concomitantly to other TZ proteins Cep290 and Cby (Figure S2A), in early G2 spermatocytes. Using expansion microscopy adapted to *Drosophila* ciliated tissues [28,29] (see methods), we show that Cfap410a-Tom decorates the growing transition zone during all steps of spermatogenesis, from early G2 spermatocytes to elongated spermatid stage (Figure 1B). Small dots of Cfap410a are observed as soon as the TZ begins to form on very short centrioles that are converting to BB. Cfap410a labeling accumulates as a discontinuous pattern along the axonemal microtubules of the TZ, while it progressively grows during spermatocyte maturation. In young elongating spermatids, Cfap410a labels the proximal region of the ciliary cap called the ring centriole. Hence, Cfap410a behaves like prototypical TZ proteins in *Drosophila* male germ cells. As the flagella grows, Cfap410a is also deposited along the axoneme in a punctate pattern (Figure S2B).

**Figure 1:**
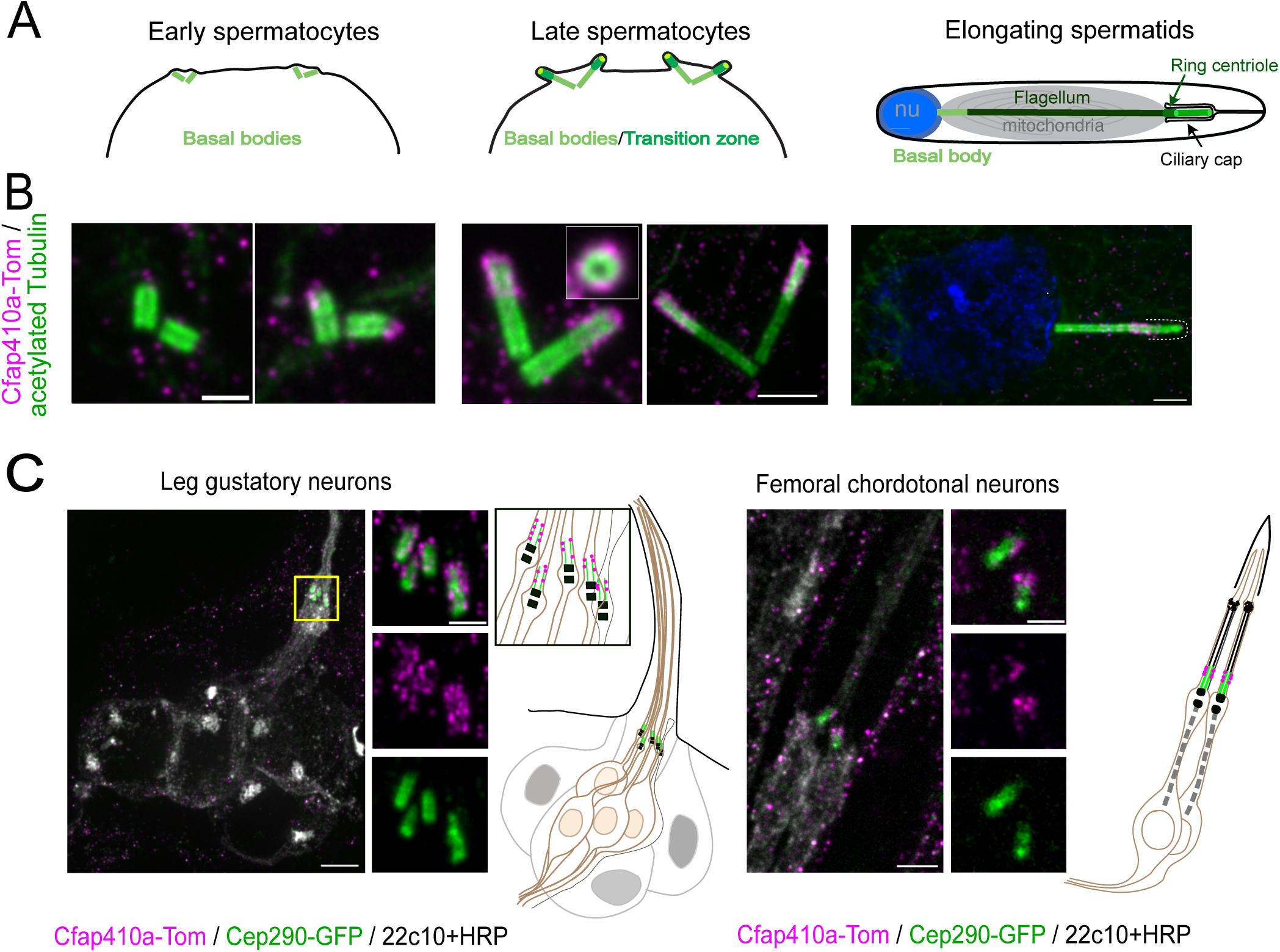
Cfap410a is located at the transition zone during spermatogenesis. (A) Schemes representing centrioles/BBs during spermatogenesis in *Drosophila* male germ cells. Two pairs of centrioles (light green) are present in each spermatocyte. During spermatocyte maturation, centrioles convert into BBs and dock to the plasma membrane, starting to build a primary cilium-like coated with TZ proteins (dark green). Both, the centriole and TZ, increase in size during spermatocyte maturation. During meiosis, the primary cilium-like is fully internalized, keeping the connection with the plasma membrane, hence forming the ciliary cap hooding the TZ/primary cilium-like. In the resulting round spermatids, BB docks to the nucleus keeping its ciliary cap inside the cell. At the base of the ciliary cap, the ring centriole (dark green) connects the base of the ciliary cap to the axoneme. During spermatid elongation, the axoneme (dark green) is assembled inside the ciliary cap and extruded inside the cytoplasm, while the ring centriole migrates along the growing flagellum end (nu = nucleus). (B) U-ExM of testes (see methods) showing Cfap410a-Tom and acetylated tubulin (centriole + TZ, green). Each stage corresponds to the scheme on top described in (A). Cfap410a-Tom starts to accumulate at the tip of the BB in young spermatocytes and fully caps the TZ in maturating spermatocytes when centriole and TZ elongate. In elongating spermatids, Cfap410a-Tom marks the base of the ciliary cap/TZ at the distal end of the spermatid cyst. Scale bars left panel = 0.5 μm mid and right panel = 1 μm. (C) U-ExM of *Drosophila* pupae legs showing the gustatory and chordotonal organs labeled with 22c10 (cell body and dendrite) and HRP antibodies (entire cell membrane including ciliary membrane) (grey). Left: gustatory organs are composed of 5 neurons (4 chemosensory neurons and one mechanosensory neuron). The TZ is labelled by Cep290-GFP (green). Enlargement of yellow inset is presented as three separated panels. In all 5 neurons, Cfap410a-Tom (magenta) surrounds the TZ labelled by Cep290 (green). Right: in femoral chordotonal neurons composed of two neurons, Cfap410a-Tom (magenta, dots on the scheme) decorates the TZ surrounding Cep290-GFP staining (green). Scale bars gustatory neurons = 2 μm, chordotonal neurons = 1 μm and both close-up = 0.5 μm.

In sensory neurons, Cfap410a was also detected at the ciliary transition zone of ciliated neurons (Figure 1C). Ciliated sensory neurons are found on diverse parts of *Drosophila* body [30]. In particular, on the legs where the two major types of ciliated sensory neurons are present: external sensory organs (gustatory neurons) and chordotonal organs. Adapting expansion microscopy to the *Drosophila* leg, we observed that Cfap410a-Tom was present in both types of neurons at the tip of their dendrites. In gustatory sensory organs, which comprise 4 gustatory neurons and one mechanosensory neuron, Cfap410a overlaps with most of the Cep290 compartment (Figure 1C, left). Cfap410a also localizes more radially outward compared to Cep290 at the TZ. In chordotonal neurons, Cfap410a staining was faint, yet still restricted to the TZ (Figure 1C, right).

In addition to this very specific signal at the TZ in both ciliated tissues, Cfap410a also shows other localizations in elongating spermatids. It is distributed along the dense body, a microtubular structure on one side of the nucleus equivalent to the human sperm manchette that shapes the nucleus during spermiogenesis (Figure S2C left: classical immunofluorescence and right: expansion microscopy). Cfap410a was also located at the leading edge of actin cones involved in sperm individualization at the end of spermatogenesis [31] (Figure S2D). These observations suggest that Cfap410a may also have functions that are independent of the TZ.

Regarding Cfap410b, the overexpression of a single copy of the Cfap410b-GFP transgene resulted in complete male sterility with abnormal elongation of the spermatids, as revealed by the dispersion of the nuclei inside the testis (Figure S3A). This precluded a precise determination of its distribution, although we observed a very high level of cytoplasmic localization (Figure S3A) and an enrichment of the protein at the primary cilium-like region only in spermatocytes (Figure S3B). These observations also indicated a strong requirement for a strict control of Cfap410b expression levels and/or a dominant negative function of the GFP-fused protein. To circumvent this limitation, we created Knock-In fly strains using CRISPR/Cas9 induced homologous recombination allowing to fuse the GFP either to the N-terminus or the C-terminus of Cfap410b (see methods). The strains are viable and fertile but none showed a reproducible localization of the protein at the TZ in spermatocytes (Figure S3C, C-terminus GFP Knock-In). The N-terminus construct did not give any signal. The C-terminus tagged strain showed a staining all along the spermatids with a specific enrichment at the distal end of the fully elongated spermatids (Figure S3D). Cfap410b was also present in the mitochondrial compartment bordering the axoneme (Figure S3E) in elongating spermatids, as confirmed by ATP5a staining. Cfap410b was also located along the dense body of the nucleus (Figure S3F). Thus, Cfap410b is enriched in non-TZ compartments in spermatids and is only present at the TZ under overexpression conditions. No Cfap410b-GFP expression could be detected in sensory neurons in none of the above strains, suggesting a faint expression in this tissue, if any.

Taken together, these findings reveal that Cfap410a and Cfap410b, each occupy unique territories within ciliated cells in the fly, with Cfap410a definitively identified as a ciliary TZ protein in *Drosophila*.

### Cfap410a is recruited downstream of Cep290, Cby and Unc at the ciliary TZ

To understand how Cfap410a is recruited at the TZ, we quantified the amount of Cfap410a-Tom in several fly mutants for TZ components (Figure 2A). The hierarchy of TZ assembly was previously described in flies [8,23,26,27,32,33] (Figure 2B). We first selected Cep290 mutant as previous studies have shown that Cep290 is a major organizer of the TZ in flies [8,23,26,27,32,33]. In the absence of Cep290, we observed that Cfap410a-Tom was almost completely absent from the TZ of centrioles, with only very few centrioles showing a Cfap410a signal (10 out of 106 counted centrioles) (Figure 2A). This indicates that Cfap410a localization is dependent on the integrity of Cep290. This later is known to be required to recruit the Dzip1-Cby-Fam92 complex [8,33]. In absence of Cby, we observed only a small but significant reduction of Cfap410a (n = 220 mutant centrioles, 440 control centrioles) (Figure 2A). This indicates that Cfap410a gets recruited at the TZ in a process that is independent of Cby, but Cby helps to stabilize Cfap410a at the TZ. Working in parallel to the Dzip1-Cby-Fam92 pathway to build the TZ, transition fiber (TF) proteins (Cep89, Fbf1 and Cep164) have been shown to be required to organize the ciliary base [28,34]. We observed that Cfap410a is not affected in *cep89^1^* mutant flies, in agreement with the previous observation that TF proteins are not required to build the TZ *per se* [28] (Figure 2A). Last, Unc has been shown to be involved both in TZ and TF formation and loss of Unc mildly affects Cfap410a recruitment (Figure 2A). Note that removal of Cfap410a in the *Cfap410a^1^* null allele (see methods) did not affect the recruitment of the following TZ components: Cep290, Fam92 or Unc (not shown).

**Figure 2:**
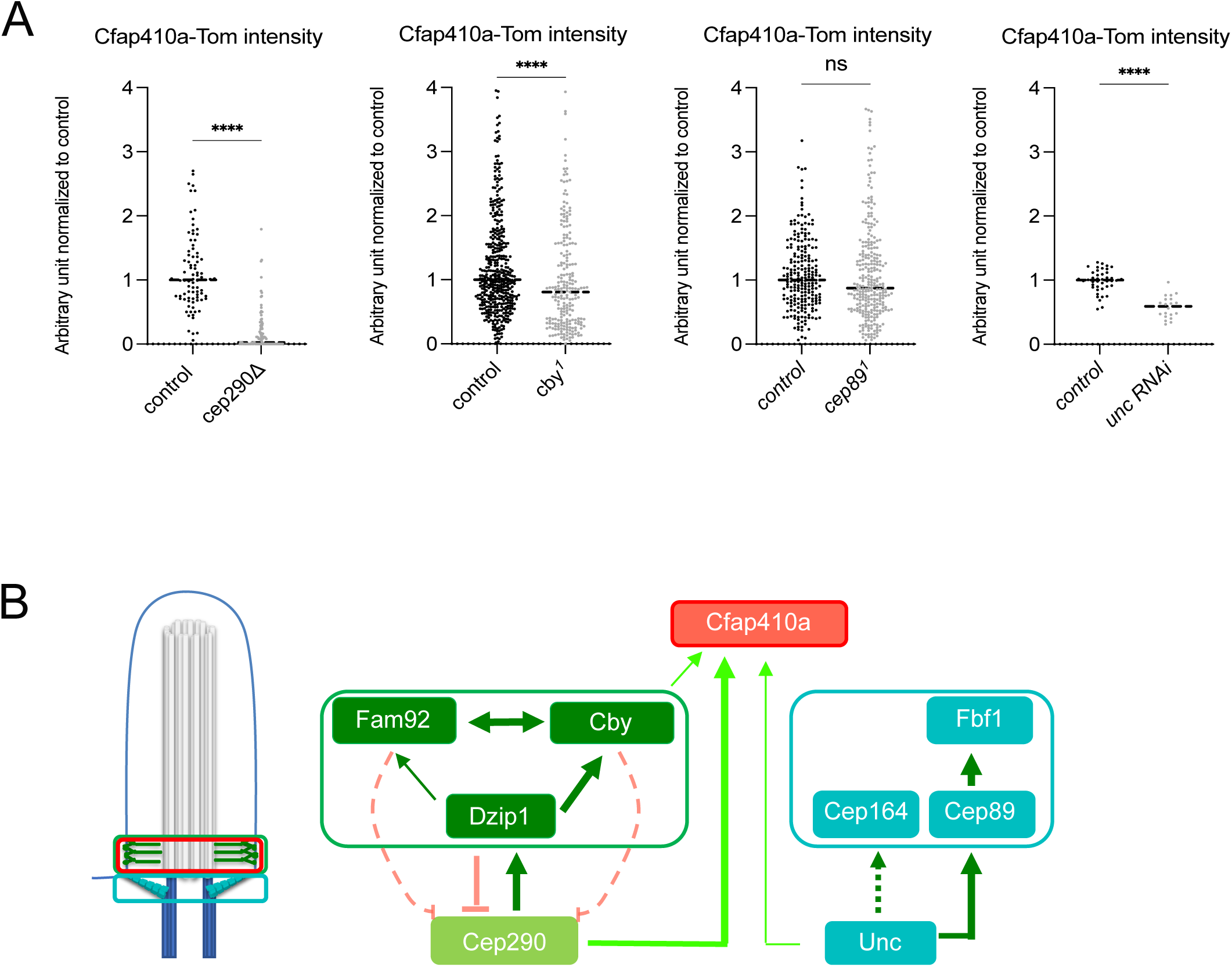
Cfap410a recruitment at the TZ depends on Cep290. (A) Cfap410a-Tom intensity was quantified in spermatocytes in control and *cep290^Δ^*, *cby^1^*, *cep89^1^* or *unc* RNAi KD mutant context. Cfap410a is almost completely absent from the TZ in absence of Cep290 and slightly but significantly reduced in absence of Cby. Removal of Cep89 does not affect Cfap410a abundancy at the TZ. Unc KD slightly but significantly affects Cfap410a abundancy. (B) Scheme of the hierarchy of TZ assembly as obtained from the literature [6,7,22–24,26,28,29,46] and positioning Cfap410a as deduced from (A).

Together, these data indicate that, in the hierarchy of *Drosophila* TZ assembly, Cfap410a lies downstream of Cep290 and is stabilized by Cby and Unc at the TZ (Figure 2B).

### Cfap410a and Cby are close proximity partners of the ciliary transition zone

We identified Cfap410a by pull-down of Cby-GFP in *Drosophila* testes (see methods). To analyze the functional relationships between Cfap410a and Cby, we first used Bimolecular Fluorescence Complementation (BiFC) assay applied to flies [35] to determine the physical proximity of the two proteins. We fused Cby to the Venus N-terminal hemi-fragment (VN) and Cfap410a to the Cerulean C-terminal fragment (CC) (Figure 3A). We used Dzip1 fused to CC as a positive control [8] and confirmed that BiFC was effective in *Drosophila* sperm cells (Figure 3B1-3). Using this assay, we observed that Cfap410a and Cby are proximity partners at the TZ of the early spermatocyte stage, when they first appear at the TZ, until the end of spermatogenesis (Figure 3B5). We also observed a BiFC interaction between Cby and Cfap410a in sensory neurons (Figure 3C). However, like for the direct Cfap410a-Tom signal, the BiFC signal was much weaker in the sensory neurons than in male germ cells. This indicates a likely reduced Cfap410a accumulation in the TZ of sensory neurons compared to male germ cells. Together, these results demonstrate that Cby and Cfap410a are close proximity partners at the TZ in *Drosophila* ciliated cells.

**Figure 3:**
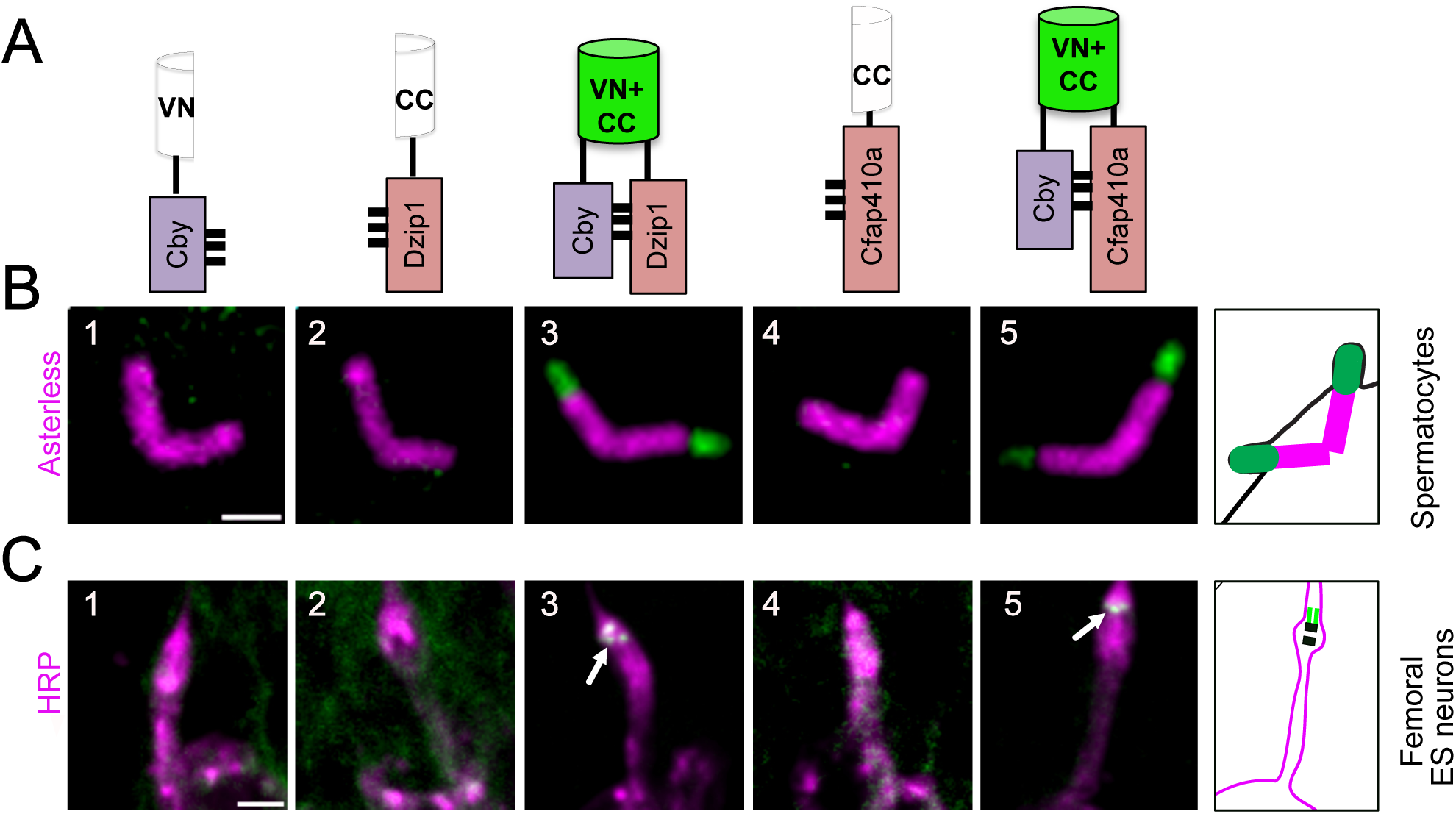
Cfap410a and Cby are close proximity partners at the transition zone. (A) Scheme representing the principle of BiFC, here with Cby fused to the N-terminal part of Venus (VN) and Dzip1 or Cfap410a fused to the C-terminal part of Cerulean (CC). Interaction between Cby and Dzip1 or Cfap410a allows the reconstitution of a functional Venus protein emitting a detectable fluorescent signal. (B) BiFC analysis of Cfap410a and Cby at the spermatocyte TZ. Flies expressing Cby- VN (1) or flies expressing Dzip1-CC (2) do not show any signal. Flies expressing both Cby-VN and Dzip1- CC (3) show a strong Venus signal at the TZ at all stages of male germ cell maturation. As well, flies expressing Cfap410a-CC (4) do not show any Venus signal during spermatogenesis whereas flies expressing both Cby-VN and Cfap410a-CC (5) show a strong Venus signal. Scale bar = 2 μm. (C) Confocal imaging of the pupal femoral ES neurons in the same conditions as in (B) allowing to distinguish a faint signal (white arrow) at the dendrite distal end in Cby-Dzip1 and Cby-Cfap410a interacting situations (3 and 5 respectively). Scale bar = 2 μm.

### Cfap410a is not essential for ciliogenesis and is not compensated by Cfap410b in fly ciliated tissues

We have created single *cfap410a^1^* or *cfap410b^1^*null mutant alleles as well as a compound *cfap410b^1^;;cfap410a^1^*mutant (see methods and Figure S4). We observed that loss of Cfap410a and Cfap410b did not affect fly development, as homozygous flies could be maintained for all three mutant strains. *cfap410a^1^* or *cfap410b^1^* and *cfap410b^1^;;cfap410a^1^* double mutant flies showed no measurable behavioral defects using conventional bang assays, suggesting that loss of either proteins or both does not adversely affect sensory cilium function that are associated to flies coordination behavior. In addition, this behavior is not aggravated by combining mutations in the two genes. These indicate that the two genes do not compensate for each other in sensory cilia assembly and function. *cfap410a^1^* or *cfap410b^1^* flies showed male fertility defects (Figure S5A-B) that were associated with weak dispersion of nuclei along spermatid cysts and slight alteration of actin cone migration (Figure S5C) highlighting possible defects in spermatid elongation and individualization [8,9,27]. The *cfap410b^1^;;cfap410a^1^* double mutant males were also hypofertile, with a delay in acquiring fertility compared to the controls (Figure S5D) but this appears only as an additive phenotype of the two single mutants (Figure S5A-B). Electron microscopy observations of *cfap410a^1^* mutant spermatids revealed weak, but significant, defects of the axoneme (Figure S5E2-3). *cfap410a^1^* mutant spermatids also exhibit disrupted association of the mitochondria and axoneme (Figure S5E5, E7 to E10), whereas *cfap410b^1^* mutant flies only show alterations of the mitochondria (Figure S6A).

Overall, *cfap410a* and *cfap410b* are not essential for sensory cilium function or for sperm flagella assembly and do not compensate for each other, in agreement with their non-overlapping localization patterns in these two tissues.

### Cfap410a and Cby cooperate functionally to build the ciliary transition zone (TZ) but with distinct requirements in different ciliated cell types

Because Cfap410a is a close proximity partner of cby, we evaluated possible genetic interactions between *cby* and *cfap410a*. We thus created the double *cby^1^,cfap410a^1^* and the *cby^1^,cfap410b^1^*mutants. Strikingly by combining *cby^1^* and *cfap410a^1^*mutants, we observed very strong genetic interactions in the two *Drosophila* ciliated tissues, that were not as strong in *cby^1^,cfap410b^1^*flies. We thus focused more in details on the characterization of the *cby^1^,cfap410a^1^*mutant.

In the testes of *cby^1^, cfap410a^1^* flies, the spermatids fail to fully elongate as observed by polyglycylated tubulin staining of the sperm tail (Figure 4A). This phenotype is reminiscent of the phenotype observed in *cby^1^*, *dilatory* double mutants [27], where we showed a total absence of recruitment of the TZ components of the MKS complex. In agreement with this observation, we found that the absence of both Cfap410a and Cby resulted in a complete lack of B9d1, as compared to the single mutants or compound heterozygotes (Figure 4B). However, the absence of both proteins did not prevent Dzip1 recruitment, which is consistent with Cfap410a acting downstream of Dzip1 (Figure 4C). Failure to form the transition zone in flies has been shown to result in the aberrant extension of axonemal microtubules in late spermatocytes [27,32]. In the *cby^1^,cfap410a^1^*double mutant, we observed aberrant axonemal extensions for almost all (95%) basal bodies, compared to around 45 % in the *cfap410a^1^,cby^1^/cby^1^*mutant (Figure 4D, E). Together, these results demonstrate that Cfap410a cooperates with Cby at the TZ to regulate the assembly of axonemal microtubules in spermatocyte and elongating spermatids. Such a functional interaction for TZ formation was not observed between Cby and Cfap410b. Indeed, although *cby^1^,cfap410b^1^* double mutants showed defects in spermatid elongation (Figure S6B), we did not observe TZ assembly defects. B9d1 was still recruited at the TZ in *cby^1^,cfap410b^1^*double mutant, as in the *cby^1^* single mutant (Figure S6C). Furthermore, we only observed a small increase in aberrant axonemal elongation in the double mutant compared to the *cby^1^* mutant (Figure S6D-F).

**Figure 4:**
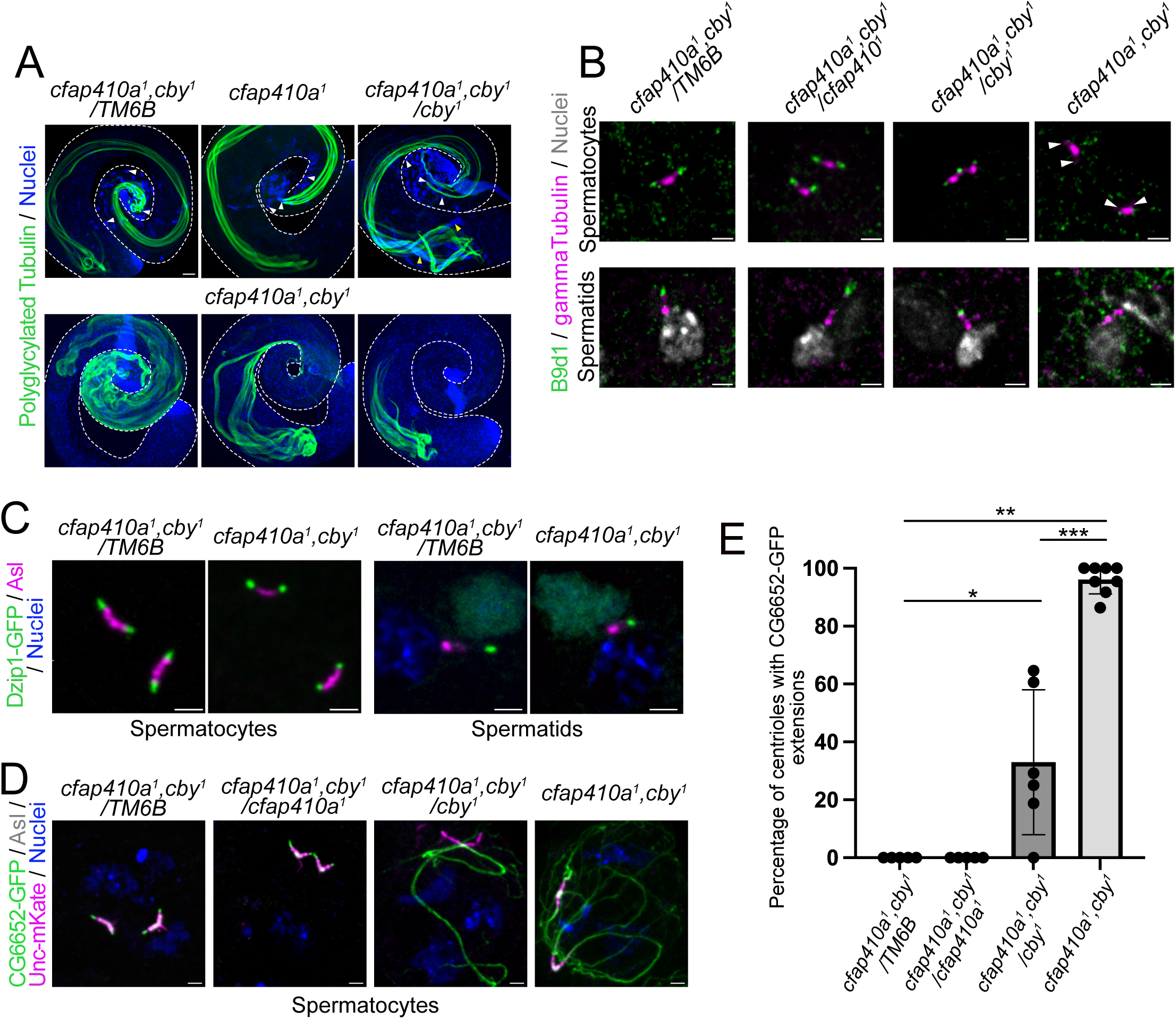
Cfap410a and Cby cooperate to build the transition zone in the *Drosophila* testes. (A) Whole mount testes showing severe defective elongation of mature spermatids (poly-glycylated tubulin labelling) in *cfap410a^1^*,*cby^1^*double mutants compared to single mutants or compound heterozygous. Moreover, nuclei bundles are completely absent in the double mutant compared to single mutant or compound heterozygous (white arrowheads). (B) The B9d1 protein is completely absent from centrioles (white arrowheads) in the double mutant compared to single mutants or compound heterozygotes at all stages of male germ cell maturation. (C) Dzip1 is still present on centrioles in *cfap410a^1^*,*cby^1^*double mutant compared to compound heterozygotes in both spermatocytes and spermatids. (D) When centriole fails to dock or when TZ integrity is compromised, abnormal elongation of centriolar microtubules is observed in spermatocytes. Centrioles (labelled with Unc-mKate) and axonemes (labelled with CG6652-GFP) in control or mutant conditions showing the aberrant elongation in spermatocytes in the double mutant compared to control. (E) Quantification of aberrant centriolar elongation defects in *cfap410a^1^*,*cby^1^* mutant conditions compared to single mutants or compound heterozygous. Mann Whitney statistical test: p < 0.05*, p < 0.01** and p < 0.001***. Scale bars: A = 50 μm, B-C and D = 2 μm

The cooperative function between Cby and Cfap410a was also observed in sensory neurons as suggested by the strong uncoordination phenotype of the flies that fail to walk when compared to single mutants in bang assays (Figure 5A). These defective behaviors were associated with an almost complete disruption of the sensory cilium of the leg femoral chordotonal organ, as revealed by almost complete loss of polyglutamylated tubulin staining in double mutant pupae (Figure 5B, C). In the antennae, chordotonal organs were analyzed by electron microscopy. In this segment, chordotonal cilia were also affected in the double mutant as indicated by a significant reduction in the number of cilia per scolopale compared to *cby^1^*or to *cfap410b^1^;;cfap410a^1^* double mutant. Nearly 40% of the scolopales were found to have fewer cilia than the control with numerous ultrastructural defects in *cfap410a^1^,cby^1^* double mutant. Several axonemes exhibited less than 9-fold microtubule (MT) doublet organisation, with 6- to 8-fold doublet arrangements and in some cases, with no accessory structures around the axonemal MTs (Figure 5D and S7). Also, most axonemes with 9 microtubule doublets lacked the inner dynein arms, and a fraction of those axonemes showed a lack of outer dynein arms and electron-dense bulges. Together, these observations indicate a strong genetic interaction between *cby* and *cfap410a* in two distinct types of chordotonal cilia in adult flies, but the severity of the defects is much stronger in the leg than in the antennae as evidenced by the complete absence of cilia in the leg compared to only 40% of missing cilia in the antennae.

**Figure 5:**
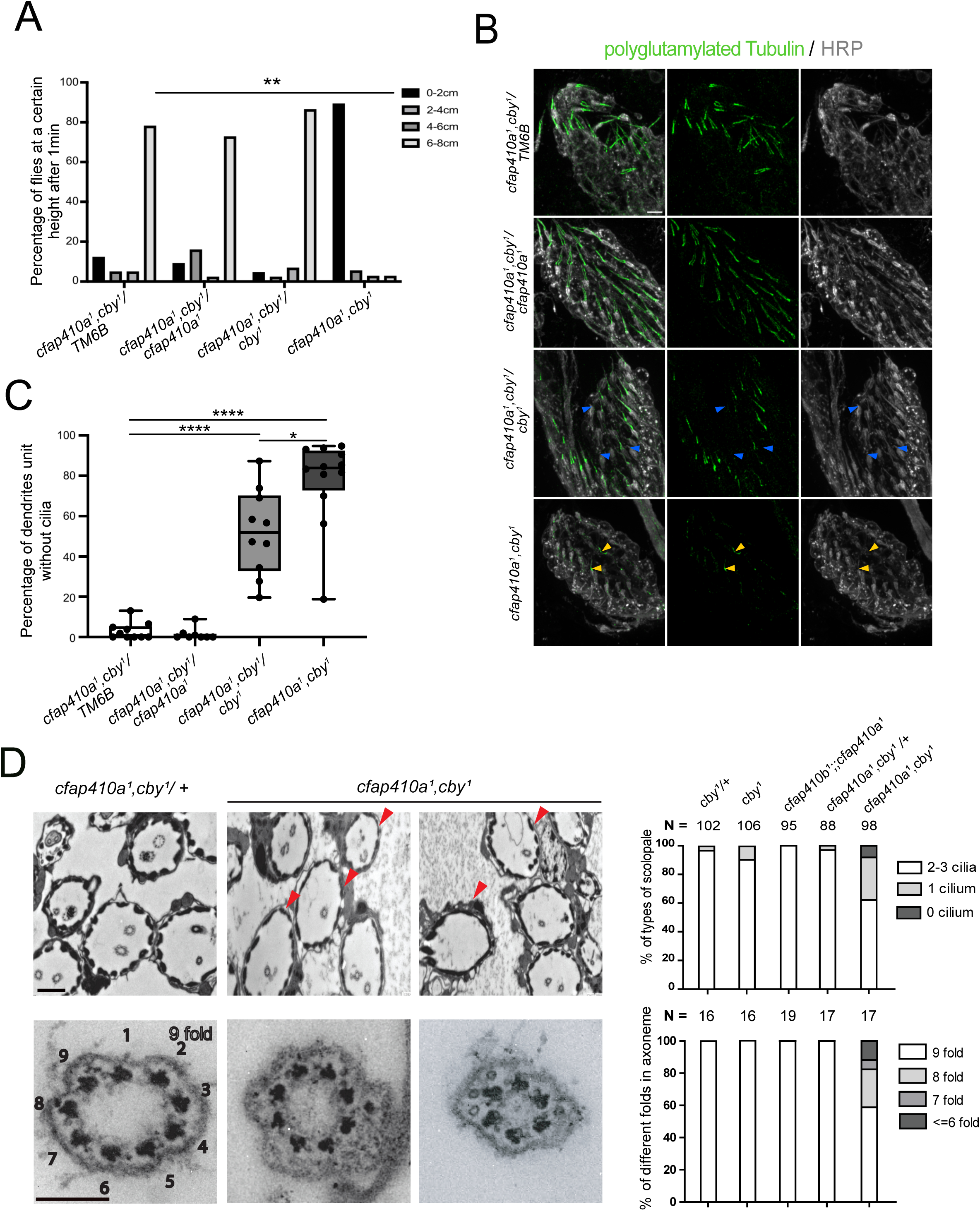
Cfap410a and Cby cooperate to build the cilium in the *Drosophila* chordotonal neurons. (A) Bang assays show strong uncoordination phenotype for the *cfap410a^1^*,*cby^1^*double mutant, with barely no flies able to move along the tube, compared to single mutants or compound heterozygous (n = 41 flies *cfap410a^1^*,*cby^1^/TM6B ;* n = 37 flies *cfap410a^1^*,*cby^1^/cfap410a^1^*,*cby^1^ ;* n = *44* flies *cfap410a^1^*,*cby^1^/cby^1^ ;* n = 44 flies *cfap410a^1^*, *cby^1^/ cfap410a^1^.* Mann Whitney statistical test: p < 0.01**. (B) Immunostaining of leg femoral chordotonal neurons show almost complete absence of the cilium at the tip of the dendrites compared to single mutants or compound heterozygous (blue arrowheads point to dendrites without cilia in the *cby^1^* mutant and yellow arrowheads point to remaining cilia in the *cfap410a^1^*, *cby^1^* double mutant). Scale bar = 5 µm. (C) Quantification of the number of cilia present in femoral chordotonal organs. Almost no cilia are detected by IF in the double *cfap410a^1^,cby^1^* (n = 10 legs *cfap410a^1^*,*cby^1^/TM6B ;* n = 12 legs *cfap410a^1^*,*cby^1^/cfap410a^1^*,*cby^1^ ;* n = 10 legs *cfap410a^1^*,*cby^1^/cby^1^ ;* n = 8 legs *cfap410a^1^*,*cby^1^/cfap410a^1^.* Statistical test Mann Whitney : p < 0.05* and p < 0.0001****. (D) Electron microscopy observations of the second antennal segment indicate a severe disorganisation of the cilia of antennal chordotonal neurons in *cfap410a^1^*,*cby^1^*double mutants compared to *cby^1^* mutant alone or *cfap410a^1^,cfap410b^1^* mutant (red arrowheads show examples of scolopodia with only one or no cilium). As indicated on the quantifications on the right, around 40% of scolopidia have no or a reduced number of cilia in the antennae in *cfap410b^1^*;;*cfap410a^1^*mutants. When cilia are present, more than 40% have an odd number of doublet microtubules as quantified on the right. See Figure S7A for more examples. Scale bar = 100 nm

We also investigated the olfactory behavior of *cfap410a^1^, cby^1^* adult flies. The olfactory cilia of the antennae’s large basiconical sensilla detect external odors, and defects in these cilia alter both electrophysiological (EAG) and behavioral olfactory responses (Figure 6A-B) [36,37,38]. *cby^1^* olfactory response has not been previously investigated. We show here that *cby^1^* mutant adults show defective olfactory behavior and reduced electrophysiological (EAG) responses compared with control flies. In contrast, the *cfap410a^1^* or *cfap410b^1^*and the double *cfap410b^1^;;cfap410a^1^* mutants do not show significantly different EAG or odor-repulsion responses compared with the control. However, *cfap410a^1^,cby^1^* homozygous double mutants are anosmic with a reduced EAG response and show a defective odor-repulsion behavior compared to *cby^1^*, confirming the genetic interaction between the two genes (Figure 6A to C). These significant behavioral defects were associated with ultrastructural defects (Figure 6D). Olfactory cilia have a characteristic outer segment (OS) made of singlet microtubules that extend from the very short cilia / transition zone present at the base of the OS and make several branches surrounded by a membrane. *cby^1^* mutant flies showed a reduced number of OS branches (by nearly 50%) and several branches were devoid of MTs compared with the control, whereas *cfap410a^1^ or cfap410b^1^* and *cfap410b^1^;;cfap410a^1^* mutants showed a normal number of outer segment branches compared with the control (Figure S7, quantification Figure 6D right). In contrast, *cfap410a^1^,cby^1^*homozygous mutant flies show a striking reduction in the total number of outer segment branches (by nearly 70%) and a significant increase in the number of branches without single MTs compared with the control. In addition, we observed disrupted transition zones (Figure 6D white arrows) in the double *cfap410a^1^,cby^1^* that were not observed in all other mutant conditions. Note that *cfap410a^1^,cby^1^* heterozygous mutant flies show normal transition zone, but the total number of outer segment branches was significantly reduced (by nearly 30%), and the number of branches without single MTs also increased significantly compared with the control (Figure S7, quantification Figure 6 right). These findings suggest that Cby and Cfap410a mutations show a genetic interaction that affects the TZ, the outer segment of olfactory cilia and the olfactory response.

**Figure 6:**
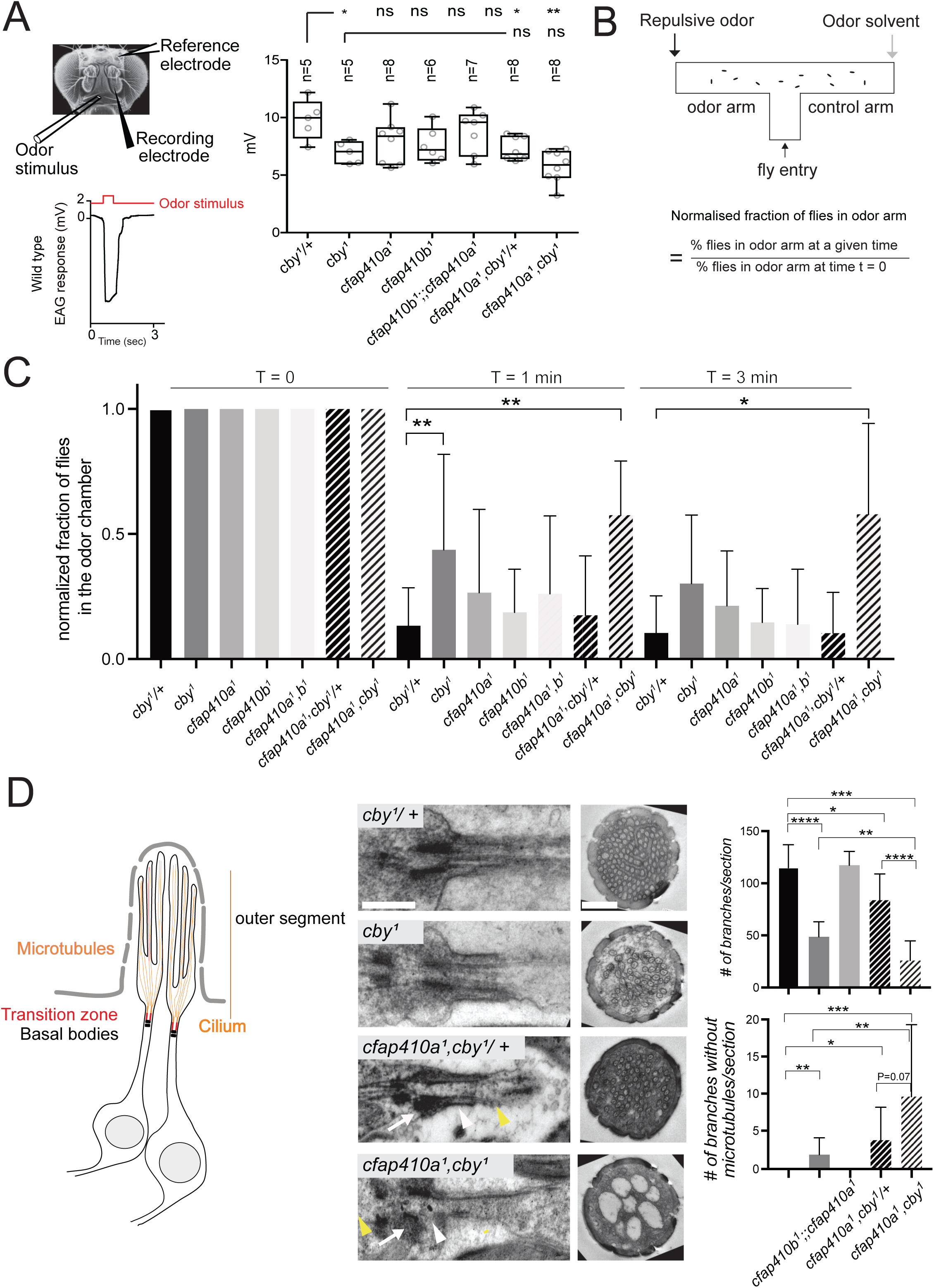
Cfap410a and Cby cooperate to build functional olfactory cilia. (A) Scheme of the electrophysiological set up to record olfactory response of *Drosophila*. For each recorded trace after olfactory stimulation, the amplitude of the peak response can be quantified (in mV) and averaged for each genetic condition. The average is plotted on the graph. Only, *cby^1^* or *cby^1^*,*cfap410a^1^* heterozygous and *cby^1^,cfap410a^1^* homozygous show a significantly reduced response, which reaches half of the control amplitude in the double mutant. (B) Scheme of the olfactory behavioral assay to evaluate olfactory response of the flies. (C) The response was quantified at several time points after odor release. Only *cby^1^* and *cby^1^,cfap410a^1^*mutant flies show a significantly altered behavioral response. (D) Electron microscopy images of the olfactory ciliated neurons in longitudinal or cross sections as indicated on the scheme on the left. In longitudinal sections, the basal body is anchored to the plasma membrane (white arrow) in all conditions but the TZ (white arrowhead) and tubular bundle (yellow arrowhead) are visible in control, *cby^1^* or *cby^1^*,*cfap410a^1^* heterozygous but strongly affected in *cby^1^,cfap410a^1^*homozygotes. Scale Bar = 500 nm. Cross section of the distal segment shows the tubular branches filled with microtubules in control conditions. In *cfap410a^1^*,*cby^1^* mutants the TZ is disrupted and the tubular branches are severely affected compared to other conditions. *cby^1^* mutants show weakly penetrant defects of the tubular branches but no apparent TZ discontinuity. Scale bar = 700 nm. See Figure S7B and C for more examples. Statistical test Mann Whitney: p < 0.05*, p < 0.01**, p < 0.001*** and p<0.0001****.

## Discussion

We show that Cfap410a, one of the two *Drosophila* CFAP410/C21orf2 orthologs, is a TZ-associated component required in different extents for the assembly or function of the three different types of cilia found in *Drosophila*. Cfap410a is not essential for organizing the TZ alone, as removing the protein causes only weak or undetectable defects in respectively the male germline and sensory neurons. However, Cfap410a works cooperatively with Cby to organize the TZ, as removing simultaneously both proteins is deleterious in all ciliated tissues in flies. Cfap410a function at the TZ is also not compensated by Cfap410b.

### Evolution of the function of Cfap410 proteins in flies

While only one member of the family is present in vertebrates, nematodes and *Trypanosoma*, *Drosophila* has two paralogs. However, our data suggest that the two proteins acquired distinct functions during evolution. Indeed, the localization of the two proteins is quite distinct, and even though both mutants exhibit fertility defects, they do not show similar phenotypes and are non-redundant, as revealed by double mutant analysis. While Cfap410a has been detected in both ciliated tissues of *Drosophila*, we were unable to detect Cfap410b in sensory neurons using various immunofluorescence conditions. Nevertheless, we cannot fully rule out the possibility that the protein may have a function in these tissues that we were unable to discern in single or *cfap410a* and *cfap410b* double mutant conditions. In the testis, Cfap410b is only observed at the TZ in overexpression conditions, but with a dominant negative effect leading to male sterility. We therefore could not conclude about a possible function of Cfap410b at the TZ, as we did for Cfap410a in sensory neurons and male germ line. Furthermore, functional interactions analysis of Cfap410b with Cby did not reveal a cooperative action at the TZ, despite the fact that both proteins cooperate for spermatid elongation (Figure S6B). In this double mutant condition we did not observe gross TZ assembly defects, as B9d1 was still recruited to the TZ as in *cby^1^*single mutant (Figure S6C).

Taken together, these observations indicate that Cfap410a and Cfap410b have evolved to exert specific and non-redundant functions in flies. It has recently been proposed that Cfap410b is required to build sensory cilia in *Drosophila*, as observed by ubiquitous induction of RNAi expression [39]. However, when inducing this RNAi in only all sensory neurons (sca-Gal4 driver line), we could not reproduce this observation, indicating that the observed phenotype (using the tub-Gal4 driver as in [39]) is likely non cellular autonomous. We also could not observe any ciliary defects in the *cfap410b* null mutant compared to the proposed complete cilia defect observed by RNAi induction [39]. These discrepancies either reveal off-target effects associated with RNAi induction or compensatory mechanisms in the null mutant, a phenomenon already described in our earlier studies [29].

### Cfap410a works cooperatively with Cby to build the TZ with tissue specific requirements

Our observations unveil Cfap410a to be a core component of TZ assembly, working downstream of Cep290 and in parallel to the previously identified Dzip1-Cby-Fam92 axis in the hierarchy of TZ assembly. In Human, mutations in *CFAP410* lead to different classes of ciliopathies and to ALS. However, the mechanisms by which CFAP410 affects cilium function is not clearly established. The protein has been shown to be part of a biochemical module that involves Nek1 and Spata7 in different cell types [40,41]. The protein has been found at centrioles/basal bodies in human or mouse cells, as well as in *Trypanosoma* [14,19,20,40], but precise localization with high-resolution imaging is missing in these systems. It has also been shown to localize along the connecting cilium of photoreceptors [14], which exhibit several features of the primary cilia TZ. *NEK1* mutations induce a reduction of CFAP410 at the centrosome in ARPE-19 cells, whereas *Spata7* mutations do not seem to affect the localization of CFAP410 at the connecting cilium of mouse photoreceptors [42]. Members of the NEK1 family are present in flies and have been shown to play numerous functions, but *Spata7* is absent from the *Drosophila* genome, suggesting that this functional module is not conserved as such throughout evolution. Our data provide another pathway by which Cfap410a links Cep290 and the *Drosophila* MKS complex. Its function is revealed only when other key TZ complexes, such as Cby, are absent. Interestingly, even in this context the severity of the observed phenotype is variable between the different ciliated cell types of *Drosophila*. In the *cfap410a^1^,cby^1^*double mutant cilia are completely missing in leg chordotonal neurons, whereas only approximately 40% of cilia are missing in chordotonal neurons of the antennae. In the testis, concomitant loss of both proteins leads to extremely severe flagella assembly defects. This emphasises that the requirement for specific complexes at the TZ differs between tissues within an organism, in addition to varying across evolution. This further highlights the need for a more comprehensive understanding of the principles governing diverse TZ assembly and composition.

### The function of CFAP proteins is likely not restricted to the TZ

Defects in mitochondria morphogenesis were observed in *cfap410a* mutants and several other mutants of transition fiber components, such as Cep89 or FBF1 [8,28,43]. It is unclear how proteins that are apparently restricted to the ciliary base in flies, like Cep89, can influence mitochondrial shaping. One hypothesis is that the TZ or ring centriole, could be directly connected to the distal end of the mitochondria and/or organizes microtubules surrounding mitochondria that are required for its elongation [44–46]. While Cfap410a-Tom is initially restricted to the TZ in spermatocytes and round spermatids, it progressively accumulates along the entire length of the spermatids as elongation starts, although the specific compartment could not be precisely identified. This compartment does not seem to be the mitochondria, based on co-staining experiment, as opposed to Cfap410b that is likely present along the mitochondria. Nevertheless, the mitochondrial shaping defect may be directly linked to this specific pool of Cfap410a. Interestingly CFAP410 was first described as a mitochondrial protein in mammals [47], whereas more recent experimental evidences show that CFAP410 exhibits a cytoplasmic and nuclear localization, and that the protein is required for mitochondrial homeostasis [48]. Together, these convergent observations suggest that, in addition to a specific role at the ciliary transition zone, CFAP410 proteins have conserved direct or indirect functions in controlling mitochondrial morphogenesis and function. In flies, this function could also be exerted by Cfap410b as we have shown that this protein is associated with mitochondria. The possible function of Cfap410b in mitochondria morphogenesis during spermiogenesis could explain the cumulative effects observed on spermatid elongation when Cby is also absent.

Cfap410 proteins in flies are also associated with the microtubules of the dense body along the nucleus, which contributes to nuclear shaping during spermiogenesis. This supports the hypothesis of a shared role of Cfap410a and Cfap410b along accessory microtubules both of the nuclei and of the mitochondria. We did not observe any specific defects of the nuclei in the mutants, even though more detailed observations may be necessary to identify possible defects.

Interestingly, Cfap410a was also detected at the base of the actin cones in spermatids. It is present in a domain that likely overlaps with the myosin VI expression domain [31]. Notably, Cep97 in flies, which caps centrioles in spermatocytes, was also shown to be associated with this domain [49]. This indicates that a conserved set of molecular components are found at the distal basal body or TZ and also at the motor end of the actin cones. Further work is required to understand the possible shared functions at these two localizations.

In conclusion, our work indicates that Cfap410a is a TZ protein that, together with Cby, functions to bridge Cep290 with the MKS components in flies. It highlights how the building of the TZ has evolved complexes that exert redundant functions, as defective TZ function can only be observed in double mutant conditions. This redundancy between TZ components could have important implications in the understanding of diseases linked to these genes. It also reveals tissue specific requirements of TZ components in *Drosophila*. Whereas Cfap410a is dispensable alone for sensory cilium formation, it is required for proper spermatid differentiation. Our work also suggests that Cfap410 proteins, in addition to having a function at the TZ, also likely impact mitochondrial homeostasis as mitochondrial abnormalities are observed in both *Cfap410* mutants. This dual function is interesting in the context of the Human pathologies associated with *CFAP410* mutations. Indeed, CFAP410 mutations have been shown to be associated with retinal and skeletal ciliopathies, but also with ALS, which affects motoneuron survival. Recently a family was described showing ALS coupled with retinitis pigmentosa. A causal point mutation c.319T>C, p.Y107H was identified in this family, which is part of the leucine rich repeat domain that is conserved from Human to *Drosophila* [50], illustrating that a single *CFAP410* mutation can lead to two diseases. In this context our observations suggest that part of CFAP410 function is mediated through its interaction with TZ components. A future challenge is to understand how variations in these interactions between tissues and patients may explain the variability of phenotypic expression of CFAP410 deficiency in Human.

## Material and methods

### *Drosophila* testis Dzip1-protein complexes purification and mass spectrometry analysis

Dzip-protein complexes were purified following a protocol adapted from [6]. Briefly, testes from ∼400 males were dissected and homogenized in 500 µl lysis buffer. The lysate was clarified by centrifugation at 14,000 × g for 20 min at 4°C. The supernatant was then incubated with GFP-TRAP beads (Chromotek) for 1h 30 min at 4°C. After several washes, protein complexes were eluted in 100 µl of 2X Laemmli Sample Buffer (Sigma) and subjected to mass spectrometry as described previously [6].

### Fly stocks and maintenance

All flies were raised on standard nutrient media between 18 and 25°C or at 29°C for RNAi experiments on a day-night cycle. The list of *Drosophila* lines used in this study can be found in Table 1.

**Table 1:**
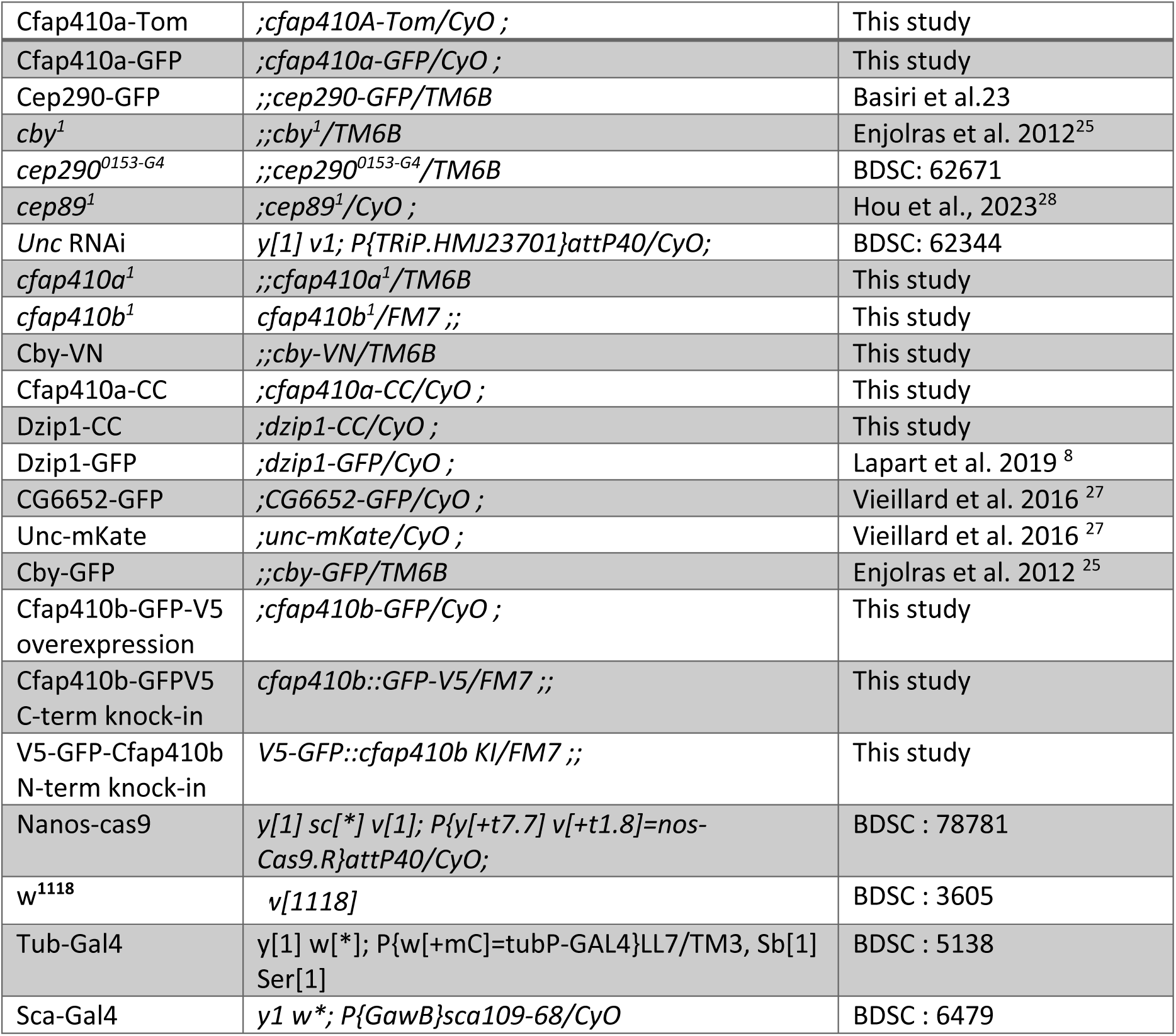
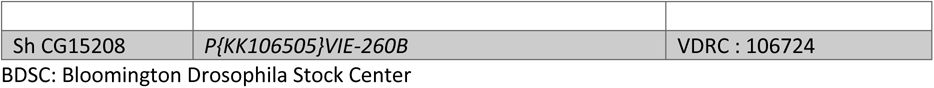
*Drosophila* stocks.

### *Drosophila* reporter gene constructs

All primer sequences are described in Table S1. All transgenic lines were obtained from BestGene Inc. Cfap410a-Tom was obtained by cloning the PCR product (primers F-CG14995Tom/R-CG14995Tom) containing 2 kb upstream regulatory sequences and the entire coding sequence (8.25 kb) of *Cfap410a/CG14995,* in frame with Tomato of pJT108 using Gibson Assembly Master Mix (New England Biolabs Inc.). For Cfap410a-GFP construct, the NotI cassette of Cfap410a-Tom, containing the entire 10,25 kb of *Cfap410a/CG14995*, was cloned in frame with GFP in NotI site of pJT61 [23]. For Cfap410b*-* GFP plasmid construct, the PCR product (F-CG15208GFP/R-CG15208GFP), including 0.87 kb upstream regulatory sequences and the entire coding sequence (1.66 kb) of *Cfap410b/CG15208*, was cloned in frame with GFP in EcoRI-NotI sites of pJT61. The resulting constructs were integrated in the 22A3 VK00037 landing site on the second chromosome by PhiC31 integrase. Cby-VN was obtained by cloning the linker-VN PCR product (F-LinkVN/R-LinkVN), obtained on template pUAST-VN, in AgeI-EagI restriction sites of Cby-Tomato [21] in replacement of the Tomato cassette. *Dzip1/CG13617-CC* was obtained by cloning the *dzip1/CG13617* 3’UTR PCR product (F-Dzip13’UTR/R-Dzip13’UTR) in XbaI restriction site of pattB-CC. The later was constructed by cloning the PCR product linker-CC (F- LinkCC/R-LinkCC) obtained on template pUAST-CC, in XhoI-XbaI restriction sites of the modified pattB containing SV40polyA sequence. pUAST-VN and pUAST-CC were a gift from S Merabet (IGFL, Lyon). The *dzip1/CG13617* upstream regulatory sequence and coding sequence (4.35 kb) was recovered by NotI-SalI digestion of *dzip1/CG13617*-GFP [6] and cloned in NotI-XhoI sites of pattB-CC-*dzip1/CG13617* 3’UTR. Cfap410a-CC was constructed as followed: *Cfap410a/CG14995* upstream regulatory sequence and coding sequence (10.25 kb) was recovered by NotI digestion of Cfap410a-Tom and cloned in NotI site of pattB-CC. *Cfap410a* 3’UTR PCR product (F-CG149953’UTR/R- CG149953’UTR) was further cloned in the XbaI site of the previous construct using Gibson Assembly Master Mix (New England Biolabs Inc.). Cby-VN was integrated in the 62E1 VK00031 landing site on the third chromosome, Dzip1-CC and Cfap410a-CC on the 53B2 VK00018 and 22A3 VK00037 landing sites of the second chromosome respectively, by PhiC31 integrase.

### Generation of cfap410a^1^, cfap410b^1^ and V5-GFP-cfap410b and cfap410b-GFP-V5 KI alleles

Both *cfap410a^1^* and *cfap410b^1^* alleles were generated by CRISPR/Cas9 induced deletion. For *Cfap410a^1^*, two gRNA1 and 2 in 5’ (5’-TCCATCATCAAGCGAATGCG-3’ and 5’- CAGTTTGGTGCAATCCTCGA-3’) and 2 gRNA3 and 4 in 3’ (5’-CGAGTAGTGATTCATCACAG-3’ and 5’- TACAAATCAGATACTTAACG-3’) of the gene were selected. gRNA1, 2 and 4 were cloned in pCFD5 vector using Gibson Assembly Master Mix (New England Biolabs Inc.) and gRNA3 oligos were phosphorylated by T4PNK (New England Biolabs Inc.) annealed and cloned in BbsI site of pCFD5. The 2 constructs were co-injected with pCFD5 expressing a validated gRNA-e (5’-GCCACAATTGTCGATCGTCA-3’) against *ebony* [45] in *nanos::cas9* embryos [44]. For *cfap410b^1^*, two gRNA in 5’ (5’- GAAGCAACTAACAGAGCACA-3’ and 5’-AAGTTGAATTTCTGTAACTG-3’) and one in 3’ (5’- GCCATTCAATGCTAACCCAG-3’) of the gene were selected and cloned in pCFD5 using Gibson Assembly Master Mix (New England Biolabs Inc.). The resulting construct was co-injected, as before, with pCFD5 expressing the validated gRNA-e against *ebony* [45] in *nanos::cas9* embryos. Deletion lines were screened with genomic PCR F-CG14995del/R-CG14995del and F-CG15208del/R-CG15208del and then confirmed by sequencing. V5-GFP-*cfap410b* and *cfap410b*-GFP-V5 KI lines were generated by CRISPR/Cas9 induced homologous direct repair. For V5-GFP-*cfap410b*, gRNA (5’- GCAACTAACAGAGCACATGG-3’) phosphorylated oligos were cloned in pCFD5 and 1,5 kb of 5’ homology arm and 1,5 kb of 3’ homology arm containing 2 silent mutations in the gRNA hybridization region, were cloned in pBSK-V5-GFP to generate the template repair. Both constructs were co-injected with pCFD5 expressing gRNA-e in *nanos::cas9.* Ebony flies were further analyzed by genomic PCR and sequencing for correct integration of V5-GFP (F-CG15208KI1/R-CG15208KI1). *cfap410b*-GFP-V5 KI line was obtained after co-injection of pCFD5 expressing gRNA (5’-GAGGACTAGAGTCTCCAGTT-3’), template repair plasmid (pBSK-GFP-V5 with 1 kb 5’ PAM mutated homology arm and 1 kb of 3’ homology arm) and pCFD5-gRNA-e. Ebony flies were then analyzed by genomic PCR and sequenced for correct integration of GFP-V5 in C-terminal of *cfap410b* CDS (F-CG152085’KI2/R-EGFP5’KI2 and F- EGFP3’KI2/R-CG152083’KI2).

### Immunofluorescence

#### 1/Whole-mount testes

Testes from adult flies or mature pupae (older than 48h) were dissected in PBS 1X, fixed 15-20 min in PBS 1X/PFA 4% at room temperature and wash 3 times in PBS 1X. After 20 min of permeabilization in PBS 1X/Triton X-100 0.3% (PBST), testes were blocked 1h in PBST/BSA 3%/sheep serum 5% (blocking buffer) at room temperature under agitation. Then they were incubated overnight at 4°C in primary antibodies (see Table 2) diluted in blocking buffer. Testes were washed 3 times 15 min in PBS 1X and incubated 1 to 2h at room temperature in secondary antibodies / dyes (see Table 3) diluted in PBS 1X. Testes were last washed 3 times 15 min in PBS 1X and were mounted in Vectashield Antifade Mounting Medium.

**Table 2:** Primary antibodies.

| Primary Antibody | Species | Source | Dilution IF | Dilution U-ExM |
| --- | --- | --- | --- | --- |
| Anti-acetylated α-Tubulin | Mouse monoclonal, IgG2b | clone 6-11B-1, T6793, Sigma-Aldrich, | | 1:500 |
| Anti-Futsch/22C10 | Mouse monoclonal, IgG1 | DSHB | 1:1000 | 1:500 |
| Anti-RFP | Mouse monoclonal, IgG2c | ChromoTek, 6G6 |  | 1:100 |
| Anti-GFP | Rabbit Polyclonal | TP401, Chemokine |  | 1:250 |
| Anti-Asl | Guinea Pig Polyclonal | Home-made | 1:50 000 |  |
| Anti-Asl | Rat | from G. Rogers | 1:75 000 |  |
| Anti-pan polyglycylated Tubulin | Mouse monoclonal IgG1 | clone AXO 49, MABS276, Sigma-Aldrich | 1:1000 |  |
| B9d1 (CG14870) | Guinea Pig | Home-made [27] | 1:1000 |  |
| Anti-γ-Tubulin | Mouse monoclonal IgG1 | clone GTU-88, T6557, Sigma-Aldrich | 1:1000 | |
| Anti-α-Tubulin | Guinea Pig polyclonal | AA345, ABCD antibodies | | 1:500 |
| Anti-β-Tubulin | Guinea Pig polyclonal | AA344, ABCD antibodies | | 1:500 |
| Anti-ATP5a | Mouse monoclonal, IgG2b | clone 15H4C4, Abcam ab14748 |  | 1:500 |

**Table 3:** Secondary antibodies and dyes.

| Antibody or dye | Reference and supplier | Dilution IF | Dilution U-ExM |
| --- | --- | --- | --- |
| Anti-mouse |  |  |  |
| Alexa fluor 488 goat anti-mouse IgG (H + L) | A11001, Invitrogen | 1:1000 |  |
| Alexa fluor 594 goat anti-mouse IgG2b | A21135, Invitrogen |  | 1:500 |
| Alexa fluor 594 goat anti-mouse IgG (H + L) | A11005, Invitrogen | 1:1000 |  |
| Anti-rabbit |  |  |  |
| Alexa fluor 488 goat anti-rabbit IgG (H + L) | A11008, Invitrogen |  | 1:500 |
| Alexa fluor 555 Goat anti-rabbit IgG (H+L) | A21428, Invitrogen | 1:1000 |  |
| Alexa fluor 647 goat anti-rabbit | A21244, Invitrogen |  | 1:500 |
| Anti-guinea pig |  |  |  |
| Alexa Fluor 594 goat anti-guinea pig IgG (H+L) | A11076, Invitrogen |  | 1:500 |
| Alexa Fluor 647-donkey anti guinea pig IgG (H+L) | 706-605-148, Jackson ImmunoResearch | 1:1000 |  |
| Anti-HRP |  |  |  |
| Fluorescein (FITC) AffiniPure® Goat Anti-Horseradish Peroxidase | 123-095-021, Jackson ImmunoResearch | 1:1000 |  |
| Alexa Fluor® 594 AffiniPure® Goat Anti-HRP | 123-585-021, Jackson ImmunoResearch |  | 1:500 |
| DyLight 649 conjugate goat affiniPure anti-HRP | 123-495-021, Jackson ImmunoResearch | 1:1000 |  |

| Dyes |  |  |  |
| --- | --- | --- | --- |
| Phalloidin - FluoProbes® 547 | FP-AZ0330 , Interchim | 1:1000 |  |
| Hoechst 33342 | 14533, Sigma-Aldrich | 20 mg/mL used at 1/1000 | 20 mg/mL used at 1/500 |

#### 2/Squashed testes

Testes from adult flies, young pupae (24h to 36h) or colored old pupae (around 90h) were dissected in PBS 1X, fixed 15 min in PBS 1X/PFA 4% at room temperature and wash 3 times in PBS 1X. Testes were placed away from each other’s on a microscopy glass slide in a drop of PBS 1X, most of the PBS was then removed and the testes were squashed with a 22 x 40 mm glass coverslip (soft pressure was applied on the testes with the back of a forceps). The slide was incubated 10 sec in liquid nitrogen, the coverslip was removed with a razor blade and the slide was incubated 3 min in pure ethanol at −20°C. The slide was washed 3 times in PBS 1X, a PAP-pen was used to make a pool around the testes which are blocked 1h in PBS 1X 0.1% Triton X-100/BSA 3%/sheep serum 5% (blocking buffer) at room temperature. Then they were incubated overnight at 4°C in primary antibodies diluted in blocking buffer. Testes were washed 3 times 15 min in PBS 1X and incubated 1 to 2h at room temperature in secondary antibodies / dyes diluted in PBS 1X. Testes were last washed 3 times 15 min in PBS 1X and were mounted in Vectashield Antifade Mounting Medium.

#### 3/Pupae’s squashed legs

White pupae (larvae that just started pupariation) were collected and kept between 24 and 36h at 25°C or 48 to 72h at 18°C. To dissect their legs, pupae were placed ventral side up on a double side tape and their case was opened using fine scissors. The pupae were removed from the case by pulling on the head and placed in a drop of PBS 1X. The top of their head and the tip of their abdomen were cut transversally and a syringe was used to inject PBS 1X inside the pupae’s envelope to remove all the internal tissues. Then, the dorsal part of the pupae envelope was cut with a scalpel to open it and the tissue was turn ventral side down to remove the excess of tissue. Finally, the tissue was flipped back ventral side up, the wings were removed with the scalpel and the legs were kept together inside their envelope. The legs were placed in a drop of PBS 1X on a microscopy glass slide. The squash and the rest of the immunostaining procedure were performed in the exact same way as for the testes (see paragraph above).

### Expansion microscopy (U-ExM) on squashed testes and legs

For both tissues, the squashed were performed in the same way as described above except that the tissue was placed on a 15 mm diameter coverslip and squashed with a 40 x 60 mm coverslip. When squashing, a small part of the 15 mm diameter coverslip is kept sticking out of the bigger coverslip to allow its retrieval. After the step in liquid nitrogen, the coverslips are placed on a metal block kept at - 20°C to slow down the warming and letting the time to remove the little coverslip pulling with tweezers on the part that sticks out.

The anchoring step was performed by incubating the 15 mm coverslip with the squashed tissue in a solution of 2% Acrylamide (AA) and 1.4% Formaldehyde (FA) in PBS 1X for 3h at 37°C. The gelation step was done by adding 4.5 µL of APS (stock solution 10% and final concentration 0.5%) in 84.5 µL of monomer solution (PBS 1X/Sodium Acrylate 19%/Acrylamide 10%/Bis-Acrylamide 0.1%/TEMED 0.5%). A drop of 28 to 35 µL of monomer is then put on a piece of parafilm on an ice-cold plate and the coverslip placed on top of it with the tissue facing the drop. The gelation was started on this ice-cold plate for 5 min before transfer for 1h at 37°C.

The gels were then punched using a 4 mm diameter biopsy punch, transferred in a six well plate and incubated with SDS 200 mM/NaCl 200 mM/Tris-HCl pH9 50 mM denaturation buffer for 5 min at room temperature. Afterwards, the gel punches were placed in 1.5 mL tube with fresh denaturation buffer for 1.5 h at 95°C. Gels were then transferred in small beakers, washed 3 x 10 min in ddH_2_O and kept overnight at 4°C in PBS 1X.

For immunostaining, gels were blocked for 2h at room temperature in PBS 1X/BSA 1% and they were incubated in primary antibodies diluted in PBS 1X/BSA 1% in a humid chamber overnight at room temperature. Gels were then washed at least 3 times 15 min in PBS 1X/Tween 0.1% incubated in secondary antibodies/dyes diluted in PBS 1X/BSA 1% in a humid chamber 5h at room temperature. Finally, gels were washed 3 times 15 min in PBS 1X/Tween 0.1% and then in PBS 1X. The day before observation, gels were expanded for 2h in 2 to 3 baths of ddH_2_O. Before observations, gels were mounted on 24 mm diameter coated coverslips (0.2 mg/mL PolyD-Lysine during 2h at 37°C or overnight at room temperature).

### Image acquisition

All images were acquired using an IX 83 inverted microscope from Olympus, equipped with a Yokagawa CSU-X1 Spinning Disk Unit, Borealis technology for illumination and Ixon3 888 EM-CCD camera from Andor. The oil immersion Plan Apochromat 60x/1.42 NA objective from Olympus was used. All images were processed and analyzed with FiJi and for UexM images the pixel scale were changed by a factor 2.

### Transmission electron microscopy

Second and third antenna segments, and testes were dissected, fixed, processed for chemical fixation, mounted and polymerized in resin following steps as previously described [25,26,38]. For antenna samples serial thin sections (60-80 nm) were cut using RMC ultramicrotome, collected on formvar- coated copper slot grids and stained with 2% uranyl acetate and Reynolds lead citrate. Samples were examined and photographed at 120 kV using a Tecnai 12 electron and Talos F20 G2 Transmission electron microscopes. Finally, the images were processed in Fiji.

### Sequence alignments

Sequences were aligned using Clustal Omega: https://www.ebi.ac.uk/jdispatcher/msa/clustalo. Alignment results were visualized using praline http://ibivu.cs.vu.nl/programs/pralinewww/

### Fertility assays

Fertility assays were performed as follow: one- or seven-days old virgin males were crossed individually with three two-days old *w^1118^* virgin females. Crosses were kept at 25°C. After five days, tubes with one or more dead flies were discarded and not included in the assay. Parental flies were removed from the tubes and all emerging offspring was counted during the next ten days. More than 30 males per condition were tested.

### Bang assay

Colored old pupae (around 90h) were collected and separated according to their genotype. Hatching flies were collecting for one to three days. The day of the assay, the flies, separated by genotype and sex, were placed by group of 10 individuals maximum in a new tube containing medium. They were left in these tubes for at least 2h to let them recover from the CO_2_ anaesthesia. Five minutes before the test, the flies were flipped in an empty tube with 2 to 8 cm graduation. The tubes were banged 3 times on the table and the behavior of the flies were recorded during 1 min. The position of the flies along the tube after 1 min was quantified. Mann Whitney statistical test was performed on the number of flies that were above the 6 cm graduation after 1 min.

### Odor repulsion assay (T-maze)

An olfaction assay was conducted to assess fly responses to the repulsive odor of benzaldehyde, using a modified T-maze assay as previously described [37]. Groups of 7-12 flies were transferred to T-mazes after cold anaesthesia and allowed to recover for 45 min. The control arm lid contained paraffin oil-soaked Whatman filter paper, while the odor arm lid contained Whatman filter paper soaked with a 10-3 dilution of benzaldehyde in paraffin oil. Concentration was standardised using 2-day-old *w^1118^*flies. Fly behavior was recorded for 15 min, with the number of flies in the odor arm counted every 15 sec from the start of odor exposure (t = 0). Each experimental day included two sets of *w^1118^ flies,* one at the beginning and one at the end of the assay, to control for variability in experimental setup and odor concentration. At least 60 flies were analyzed per genotype. Data was plotted in GraphPad Prism. Mann Whitney statistical test was performed.

### Electrophysiology / Electroantennogram (EAG) recording

Electrophysiological recordings followed a modified protocol from previous studies [36,38] using an in-house-built setup. EAGs were obtained using in-house Ag/AgCl electrodes in NaCl-filled glass capillaries. The ground electrode was inserted into the head capsule, and the recording electrode was positioned on the large basiconic sensilla of the third antennal segment. For each recording, 10-4 ethyl acetate diluted in paraffin oil was delivered near the recording electrode for 500 ms using a LabView-controlled olfactometer. Response traces were captured with LabView software and normalized with a 20-point moving average. Normalized peaks were measured in Microsoft Excel, and the average of three recordings was plotted in GraphPad Prism. Recordings were performed on at least six flies per genotype.

## Quantification and Statistics

**Quantification of the intensity of TZ proteins in spermatocytes** were done as described in [25] on at least 80 centrioles and 2 males. **Quantification of aberrant CG6652-GFP extension in the double mutants:** the quantifications were done on cells that were in meiosis either in the first division round or the second. The number of centrioles showing aberrant extension of CG6652-GFP were quantified in all the cells in meiosis in each picture and the percentages were made per testis. Quantifications were done using FIJI cell counter plugin. **Quantification of cilia lost in pupae femoral chordotonal neurons:** in this tissue, two chordotonal neurons project their dendrites in the same direction really closely to each other. Since it is sometimes difficult to distinguish the two dendrites and their single cilium, we quantified the presence of at least one cilium in a chordotonal unit (two neurons). The percentages were made per legs. Only the absence or presence of cilia was quantified, not their length. Quantifications were done using FIJI cell counter plugin. **Quantification of ciliary axoneme formation and ultrastructural defects in adult second segment chordotonal neurons:** axonemés presence, fold symmetry of microtubule doublets and presence of different non-microtubule features were quantified using Fiji from TEM cross-sectional images of proximal region of the chordotonal cilia found in the second antennal segment. These properties were compared with those of respective controls. **Quantification of EAG response (in mV) measured from third antenna of adult flies in response to odor:** the response traces exported from LabView were smoothened using 20-point averaging and the voltage peaks were measured after zero correction in Microsoft Excel. At least 5 flies of each genotype were analyzed and average of 3 recordings from a given antenna was plotted with control for comparison of electrophysiological response. **Quantification of various ultrastructural features of the outer segment of the olfactory basiconic cilia:** branches and microtubule singlets were quantified using Fiji from TEM cross-sectional images of outer segments with a diameter of 2 µm to ensure consistency in the region analyzed. Branches with and without microtubule singlets were also counted to compare the defects in branching pattern with respective controls. GraphPad Prism was used for statistical analyzes and the creation of graphs.

## Acknowledgments

AB was supported by a doctoral fellowship from the Université Claude Bernard Lyon-1 and from the FRM. JAL was supported by the FRM (DEQ20131029168. E.F. was supported by a fellowship from the Ligue National Contre le Cancer. MD and PP are supported by the NCBS-Tata Institute for Fundamental Research (TIFR), CSIR (Council of Scientific & Industrial Research, India) Fellowships/Grants/Contracts. HAD is supported by Centre Franco-Indien pour la Promotion de la Recherche Avancée (CEFIPRA) grant. This work was supported by the ANR (ANR-22-CE13-0014 BBDIV to BD); NCBS-TIFR-DAE (an intramural grant from Tata Institute of Fundamental Research, Department of Atomic Energy (TIFR-DAE) to SCJ), Department of Biotechnology, Government of India (BT/PR53868/BMS/85/255/2024 to SCJ) and Centre Franco-Indien pour la Promotion de la Recherche Avancée (CEFIPRA) (6903-1 to SCJ and BD), Ministry of Earth Science, Government of India (MoES/PAMC/Dom/66/2023(E-14508) to SCJ). The authors acknowledge the contribution of SFR Santé Lyon-Est (UAR3453 CNRS, US7 Inserm, UCBL) facility: CIQLE (a LyMIC member) and of the NCBS Central Imaging & FACS Facility, and NCBS Electron Microscopy Facility and the NCBS fly facility. We thank Julien Perrichet for technical assistance and fly husbandry. We than Jean-Luc Duteyrat and Elisabeth Cortier for technical assistance. We acknowledge Shriya for help in some olfactory behavior experiments, Amit Morarka for establishing electrophysiology set-up assembly. We thank SCJ’s Lab (Organelle Biology Lab) members for reviewing the manuscript and providing helpful discussions on the manuscript. Mass spectrometry experiments were carried out using the facilities of the Montpellier Proteomics Platform (PPM-PP2I, BioCampus Montpellier), and were partially supported by the French National Research Agency ProFI fundings (Proteomics French Infrastructure, ANR-24-INBS-0015). Fly stocks were obtained from the Bloomington Drosophila Stock Center (NIH P40OD018537) and the Vienna Drosophila Resource Center (VDRC).

## Contributions

AB: Investigation; Methodology; Formal analysis; Writing—original draft; Writing—review and editing. JV: Investigation; Methodology; Formal analysis; Writing—original draft; Writing—review and editing. PP; Investigation; Methodology; Formal analysis. Writing—review and editing. MD: Investigation; Methodology; Formal analysis. Writing—review and editing; EF: Investigation. JAL: Investigation. SDF: investigation. HAD: Investigation. VM: resources. JT: Conceptualisation; Methodology; Investigation; Formal analysis; Supervision; Writing—original draft; Writing—review and editing. SCJ: Conceptualisation; Supervision; Funding acquisition; Writing—original draft; Writing—review and editing. BD: Conceptualisation; Supervision; Funding acquisition; Writing—original draft; Writing— review and editing.

---

**Figure S1:**
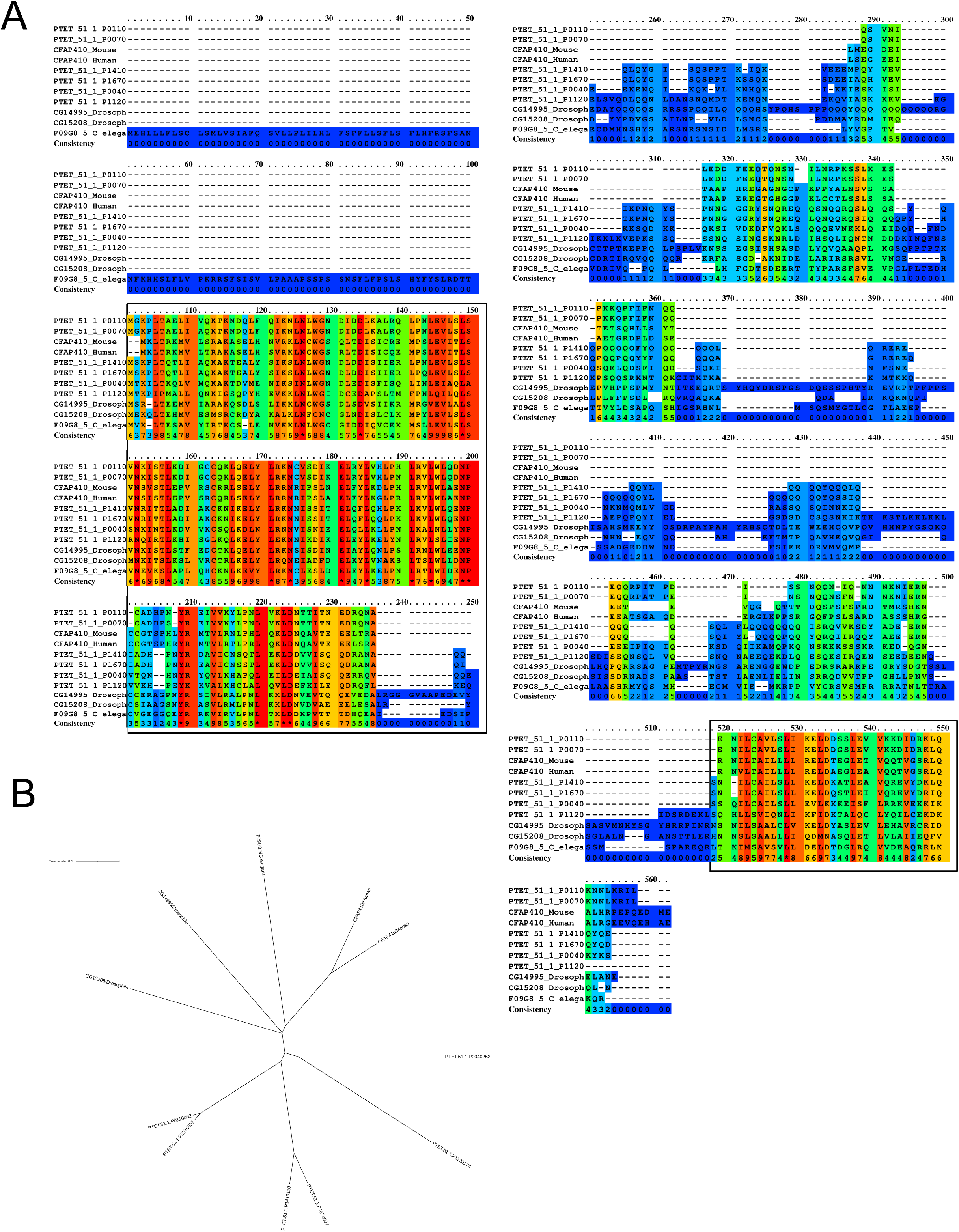
Protein sequence comparison of Cfap410a and Cfap410b. (A) Sequences from *P. tetraurelia, C. elegans, D. melanogaster, M. musculus, H. sapiens* were aligned using ClustalW. Amino acid conservation scoring is performed by PRALINE [47]. Zero corresponds to the least conserved alignment position (blue) and 10 to the most conserved alignment position (red). A Leucine Rich domain is highly conserved in N-ter of the protein and a second conserved domain is found at the C-ter (black boxes). (B) Phylogenetic tree of CFAP410 proteins, using phylogenetic tree generation methods from the ClustalW2 package (EMBL’s European Bioinformatics Institute).

**Figure S2:**
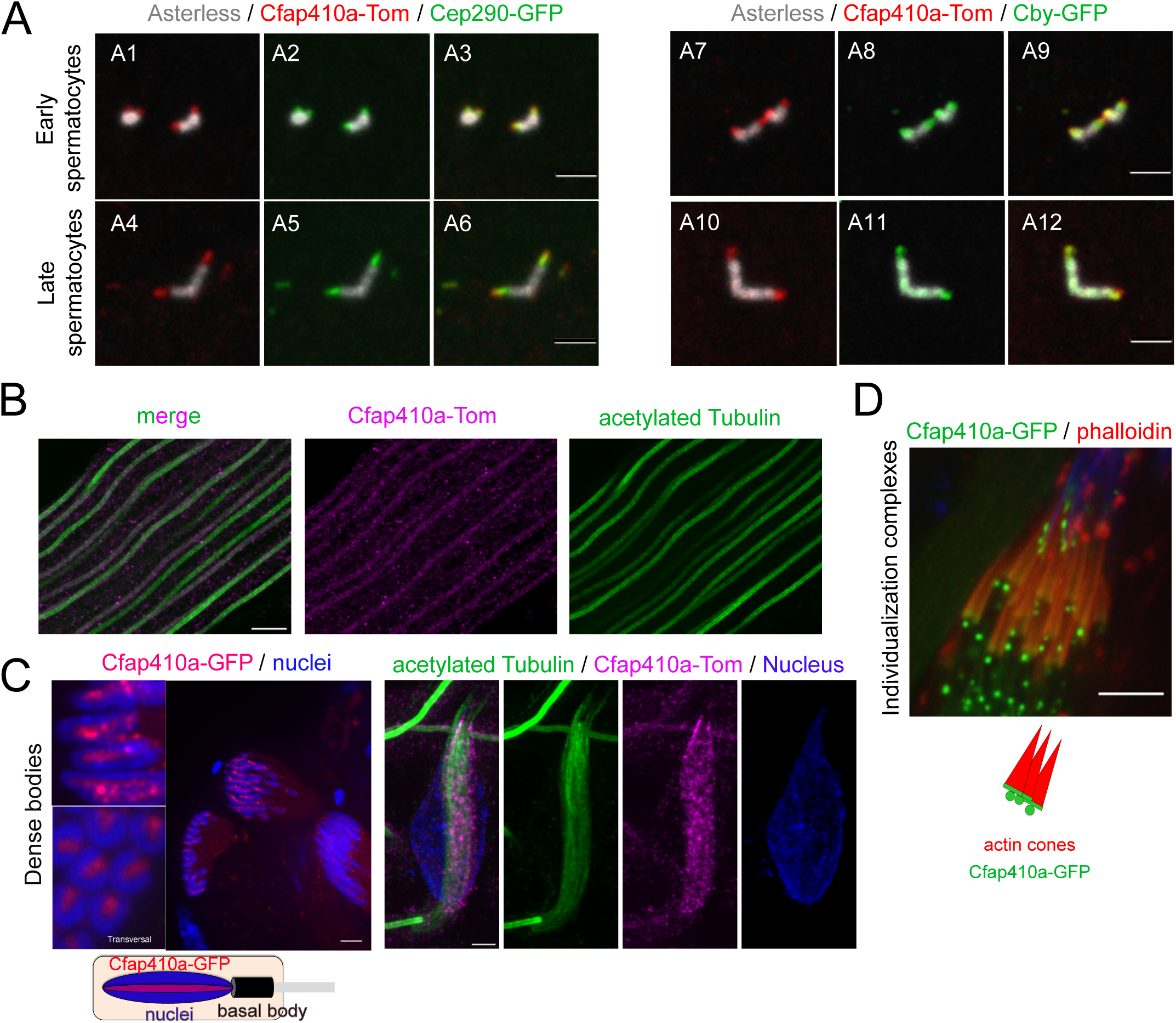
Cfap410a expression during the different stages of spermatogenesis. (A) Cfap410a-Tom and Cep290-GFP (A1-6) or Cby-GFP (A7-12) appear simultaneously at the TZ. In young spermatocytes (A1-3, A7-9), the two centrioles in each pair are very short, and Cfap410a and Cep290 or cby are always present together at the distal end of the centrioles. In later stages (A4-6, A10-12), centrioles elongate and Cfap410a-Tom is present with Cep290 or Cby at the TZ. Scale bars = 2 μm. (B) Expansion microscopy observations of elongated spermatids labeled for acetylated tubulin (green) and Cfap410a-Tom showing the punctate Cfap410a pattern along the axoneme. Scale bar = 2 μm. (C) Cfap410a is located at the dense body as observed on longitudinal and transversal views in enlargement boxes on whole mount testes. Scale bar = 10 μm. Right panels: expansion microscopy observation of the dense body (labeled by acetylated tubulin, green) along the nucleus (blue) shows the punctate pattern of Cfap410a-Tom (magenta). Scale bar = 1 μm. (D) Cfap410a-GFP is observed at the base of the actin cones (phalloidin, red). Scale bar = 10 μm.

**Figure S3:**
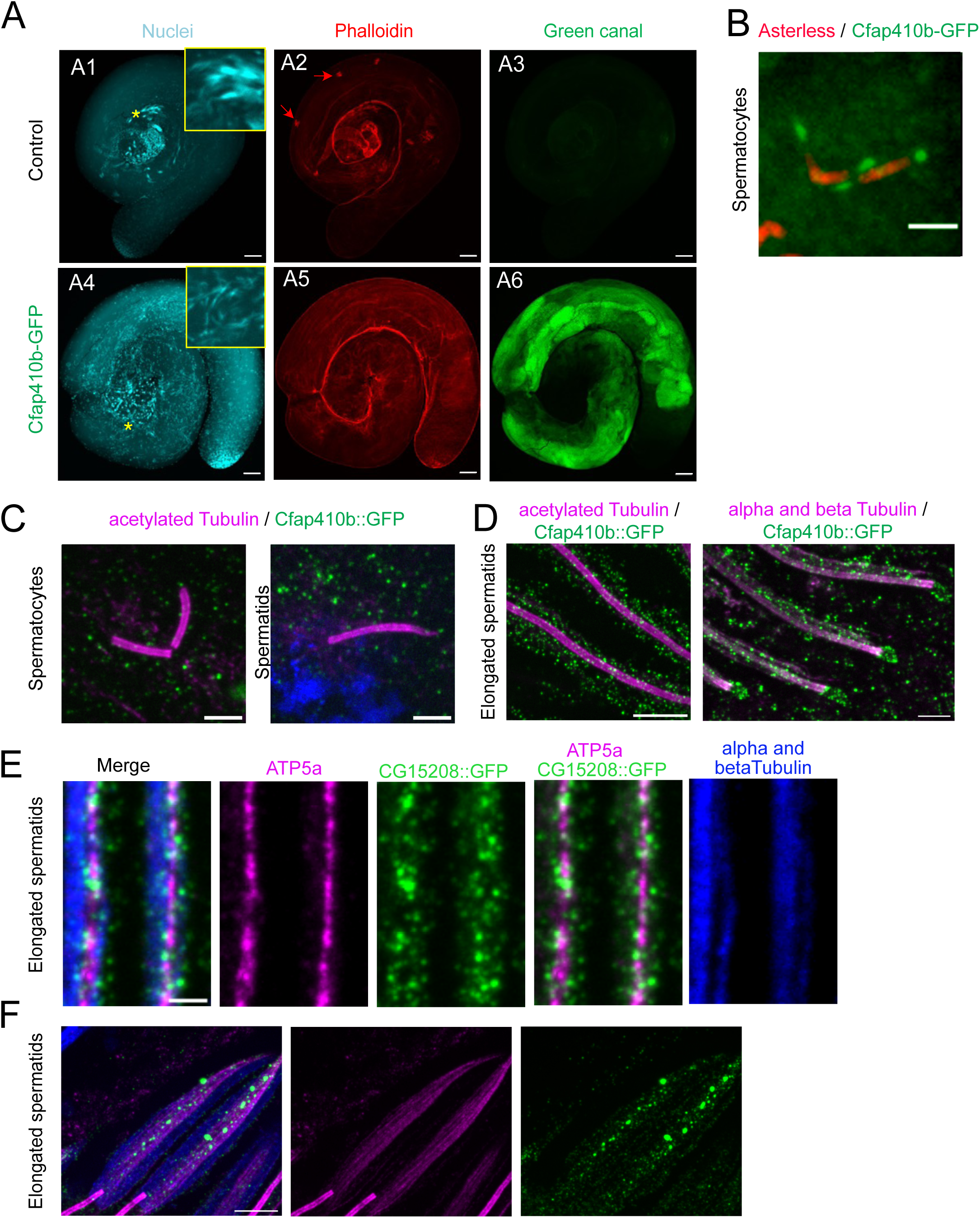
Localization of Cfap410b during spermatogenesis. (A,B) Whole mount *Drosophila* testes from fly strain overexpressing Cfap410b-GFP. (A) A strong cytoplasmic GFP signal is observed from spermatocytes to elongated spermatids compared to control non-transgenic strains (A6 compared to A3). In the Cfap410b-GFP overexpressing strain, the cysts are disorganized, with nuclei dispersed along them compared to control (A4 compared to A1), and there is an absence of actin cones (red arrows) labelled with phalloidin (A5 compared to A2). (B) Cfap410b- GFP signal is observed at the TZ in spermatocytes above a strong uniform cytoplasmic signal. (C to F) Expansion microscopy of squashed testes from Cfap410b::GFP C-term knock-in. (C) No GFP signal could be detected at the TZ both in spermatocytes and early spermatids. Scale bars = 1 μm. (D) Cfap410b::GFP is enriched around the axoneme and at the distal end of mature spermatids. Scale bars = 2 μm. (E) Cfap410b::GFP is present all along the spermatids, associated with the mitochondrial compartment labelled with ATP5a. Scale bar = 500 nm. (F) Cfap410b::GFP is present in a dotty pattern along the microtubules of the dense body in elongated spermatids. Scale bars = 2 μm.

**Figure S4:**
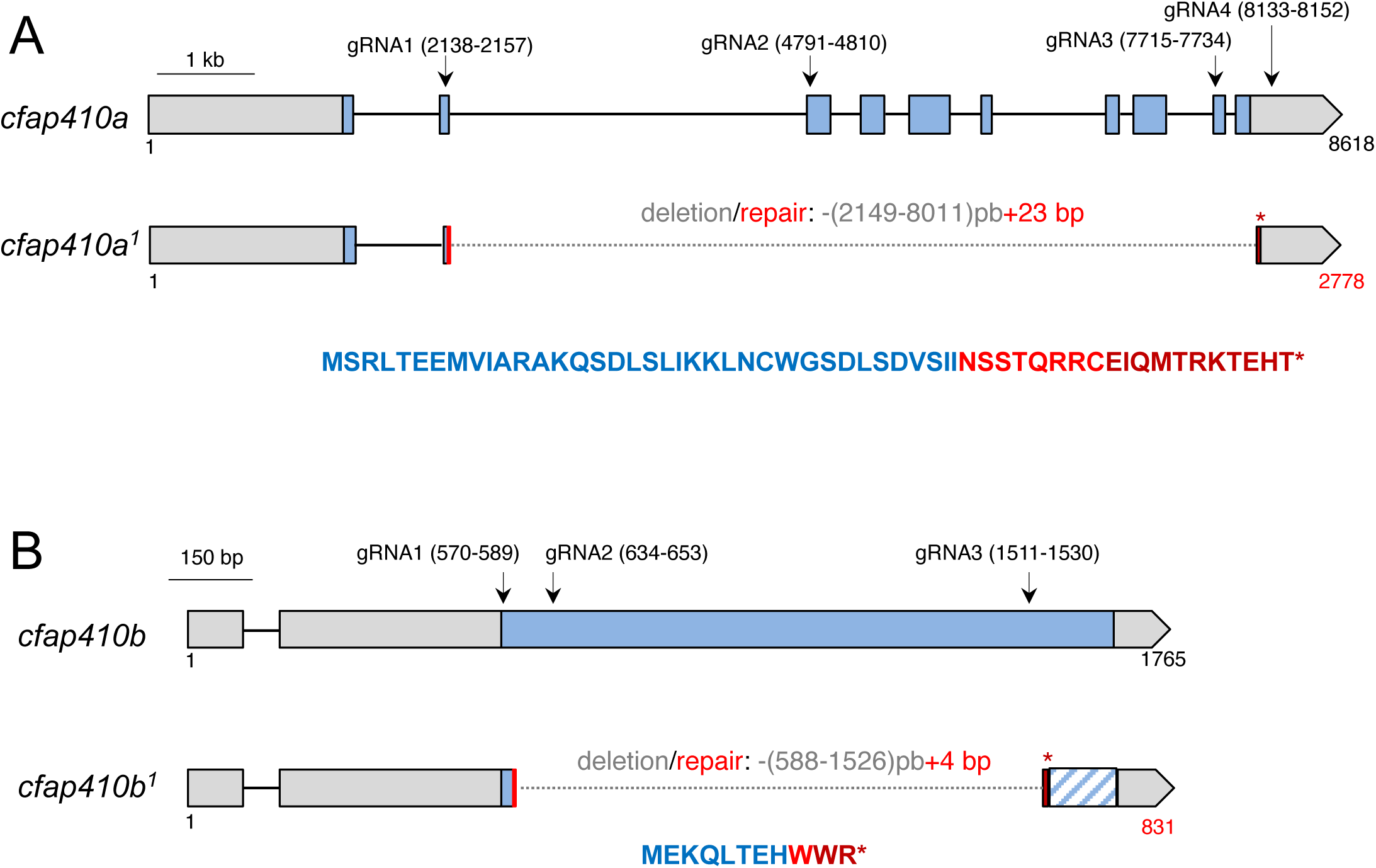
Schemes of *Cfap410a^1^* and *Cfap410b^1^* mutant alleles. (A) Scheme of *cfap410a^1^* null allele using CRISPR/Cas9 system that generated the deletion of the majority of the coding sequence. The remaining sequence can potentially produce a small chimeric protein of 56 aa comprising only the first 37 wild type aa compared to the 454 aa of the wild type protein. (B) Scheme of *cfap410b^1^* null allele. Deletion by CRISPR/Cas9 included the majority of the coding sequence. Only an 11 aa peptide can potentially be produced by the remaining coding sequence compared to the 365 aa composing the wild type protein.

**Figure S5:**
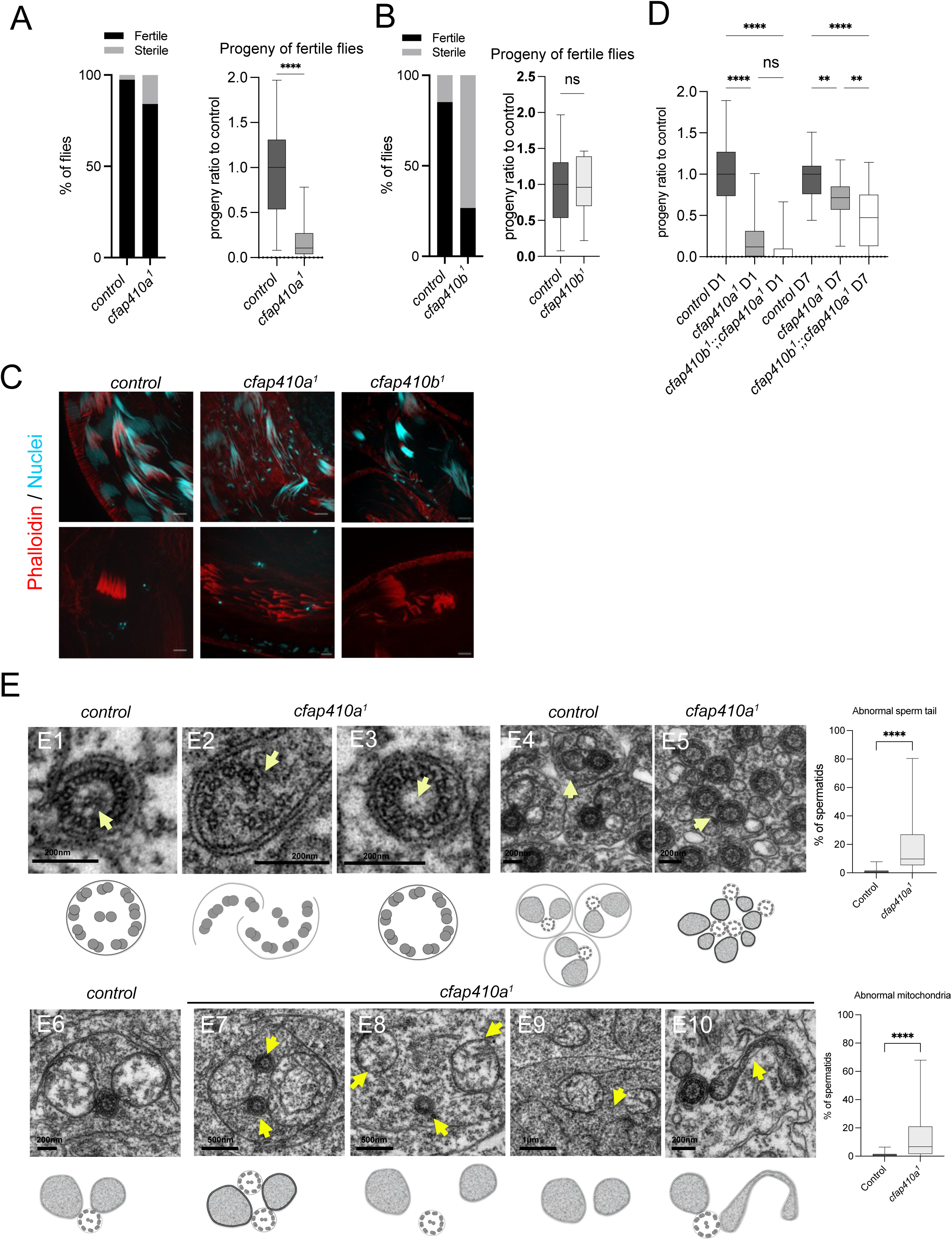
Cfap410a is required for male fertility and sperm tail formation and is not compensated by Cfap410b. (A-B) Fertility test of *cfap410a^1^* and *cfap410b^1^* single mutant. (A) Histograms representing the percentage of males with progeny in individual crosses. Left: A small but significant proportion of *cfap410a^1^*mutant males do not exhibit any progeny. Right: in the remaining flies, *cfap410a^1^*mutant males show a reduction in the average number of progenies. (B) Histograms representing the percentage of males with progeny in individual crosses. Left: in single *cfap410b^1^* mutants, 70% of males are completely sterile compared to 16.6% in the control condition (n = 36 out of 52 tested flies). Right: histogram representing the number of progenies for each fertile flies scored in left (n = 43 for control and n = 16 for *cfap410b^1^* flies tested). No significant differences could be observed between *cfap410b^1^* and control flies. (C) Immunofluorescence of whole mount testes showing weakly penetrant de- clustering of nuclei and actin cones in both *cfap410a^1^* and *cfap410b^1^* single mutant. (D) Histogram representing the number of progenies for each fertile flies in *cfap410a^1^* single mutants or *cfap410b^1^*;;*cfap410a^1^* double mutants at day 1 and day 7. No significant difference was found between males at day 1 between *cfap410a^1^* or *cfap410a^1^;cfap410b^1^* double mutants, with a slight significant reduction at day 7 corresponding to an additive effect of the two mutations. Student T-test: p-value = 0.9620>0.05. ****: p<0.0001; **: p<0.01; ns: p>0.05. (E) Electron microscopy observation of spermatid tails in control or *cfap410a*^1^ mutant testis. (E1) In control spermatids, the axoneme shows a 9+2 architecture. The arrow points to the central pair. (E2) In rare *cfap410a^1^* mutant spermatids, axonemes can be broken (arrow, E2) or the central pair is missing (arrow, E3) (8 out of 213 counted spermatids), (E4-E5) larger views of young spermatid cysts. (E4) In control testes, each cell is surrounded by its own membrane (arrow), whereas mutant cysts (E5), show fused spermatids as revealed by apposed axonemes (arrow). Note the aberrant size of mitochondria and odd mitochondria to axoneme ratio. The histogram shows the % of abnormal spermatids observed per cysts (n = 19 cysts for mutant, 17 cysts for control). Student T-test p-value: 4.75784E-8<0.001. (E6- E10) Strikingly, *cfap410a^1^* leads to rare severe mitochondrial defects during cyst elongation. (E6) In control testes, the major and the minor mitochondria are apposed to the axoneme. (E7) In mutant condition, two axonemes (arrows) can be associated with the same mitochondria. (E8-E9) The association between the axoneme and the mitochondria can also be disrupted (arrows), resulting for the most drastic cases (E9) in isolated mitochondria (E10, arrows). The morphology of the mitochondria can also be altered in *cfap410a^1^*mutants leading to “snake shape” sections (arrows) instead of round ones. The histogram shows the percentage of abnormal mitochondria observed among total observed cysts. All mitochondrial defects were pooled and quantified. Student T-test p-value: 2.74013E-5<0.001.

**Figure S6:**
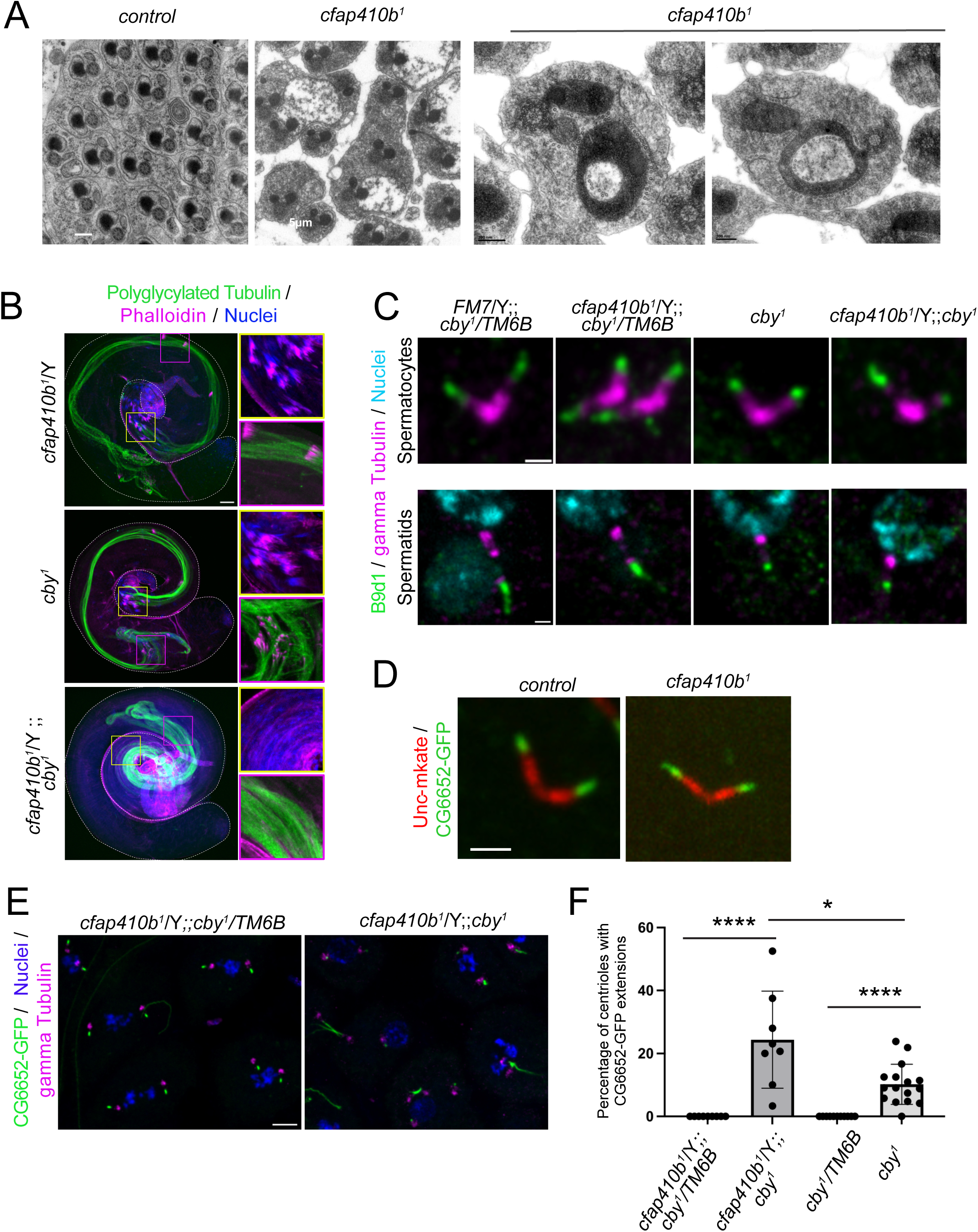
*Cfap410b^1^ mutants* show abnormal spermatid mitochondria morphogenesis and cooperate with Cby in spermatid elongation but not TZ assembly. (A) Electron microscopy observations of *cfap410b^1^* mutant spermatids. Whereas no axonemal defects can be observed, misshapen mitochondria are observed in mutant spermatids. (B) Whole mount testes showing defective elongation of mature spermatids labelled with axo-49 (polyglycylated tubulin) in *cfap410b^1^;;cby^1^* double mutant compared to single mutants. Scale bar = 50 μm. (C) B9d1 is still present in *cfap410b^1^;;cby^1^*double mutants compared to its complete absence in *cfap410a^1^,cby^1^*(Figure 4B). Scale bar = 1 μm. (D) No aberrant axonemal extension is observed in *cfap410b^1^* mutant spermatocytes. (E) CG6652-GFP recruitment is only slightly affected in *cfap410b^1^*,*cby*^1^ double mutants compared to *cby^1^* mutant alone. Scale bar = 5 μm. (F) quantifications of (E) n = 11 testes *cby^1^/TM6B*; n = 16 testes *cby^1^/cby^1^*; n = 9 testes *cfap410b;;cby^1^/TM6B* and n = 8 testes *cfap410b;;cby^1^/cby^1^*. Statistical test Mann Whitney: p < 0.05* and p < 0.0001****.

**Figure S7:**
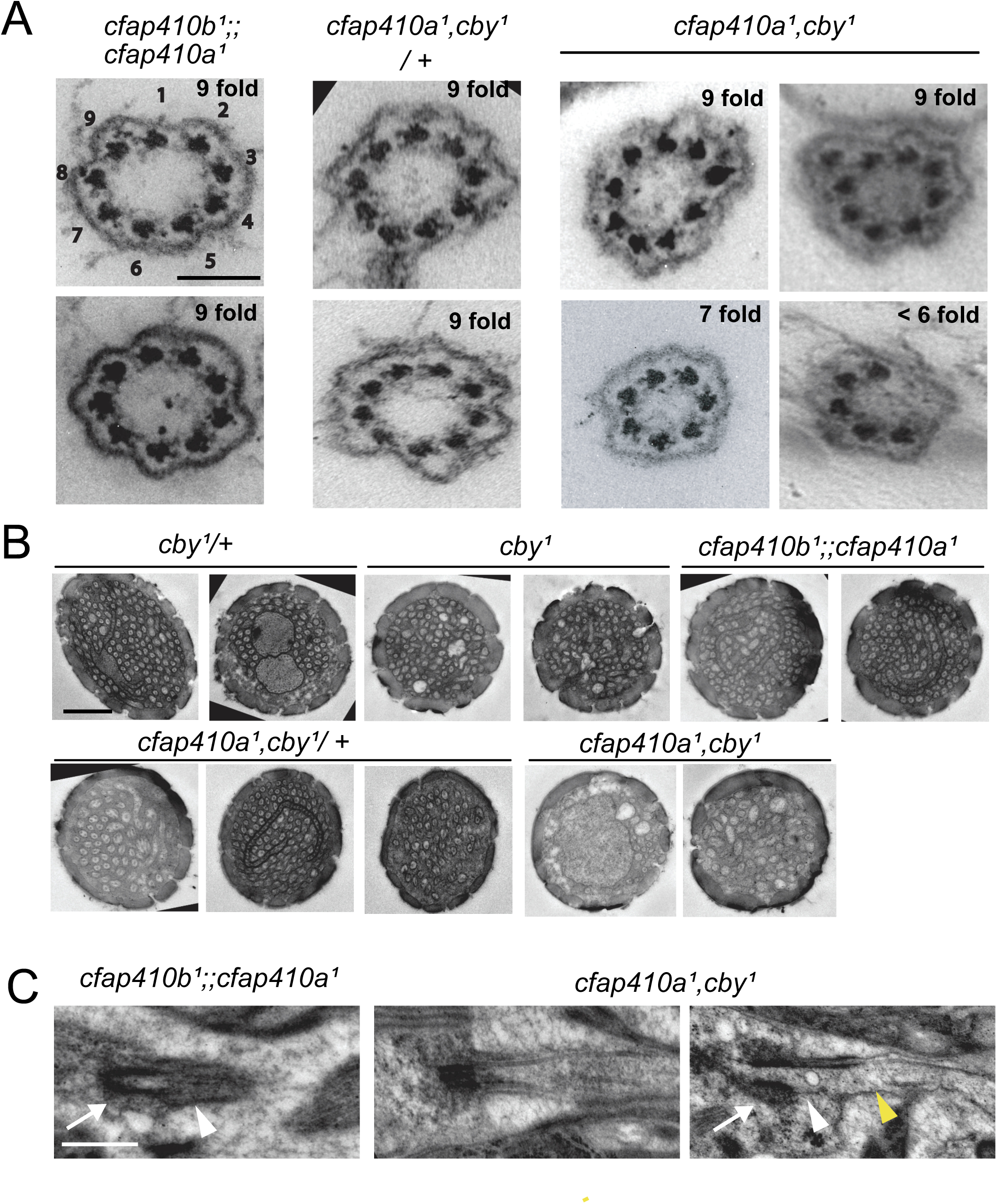
Representative electron microscopy images (relative to quantifications of Figure 5 and Figure 6) (A) Cross sections of cilia from chordotonal organs of the second antennal segment. Note the various odd numbers of doublet microtubules in *cfap410a^1^,cby^1^* double mutant compared to control heterozygous or to *cfap410a* and *b* double mutant. Scale bar = 100 nm. (B) Cross sections of olfactory sensilla of the third antennal segment representative of the different genotypes. Scale bar= 700 nm. (C) Longitudinal sections of olfactory sensory cilia in *cfap410a^1^,cby^1^* double mutant compared to *cfap410b¹;;cfap410a¹.* Scale Bar = 500 nm. On the right panel, the basal body (white arrow) is anchored to the plasma membrane but the TZ is discontinuous (white arrowhead) and tubular bundle are malformed (yellow arrowhead).

**Table S1:**
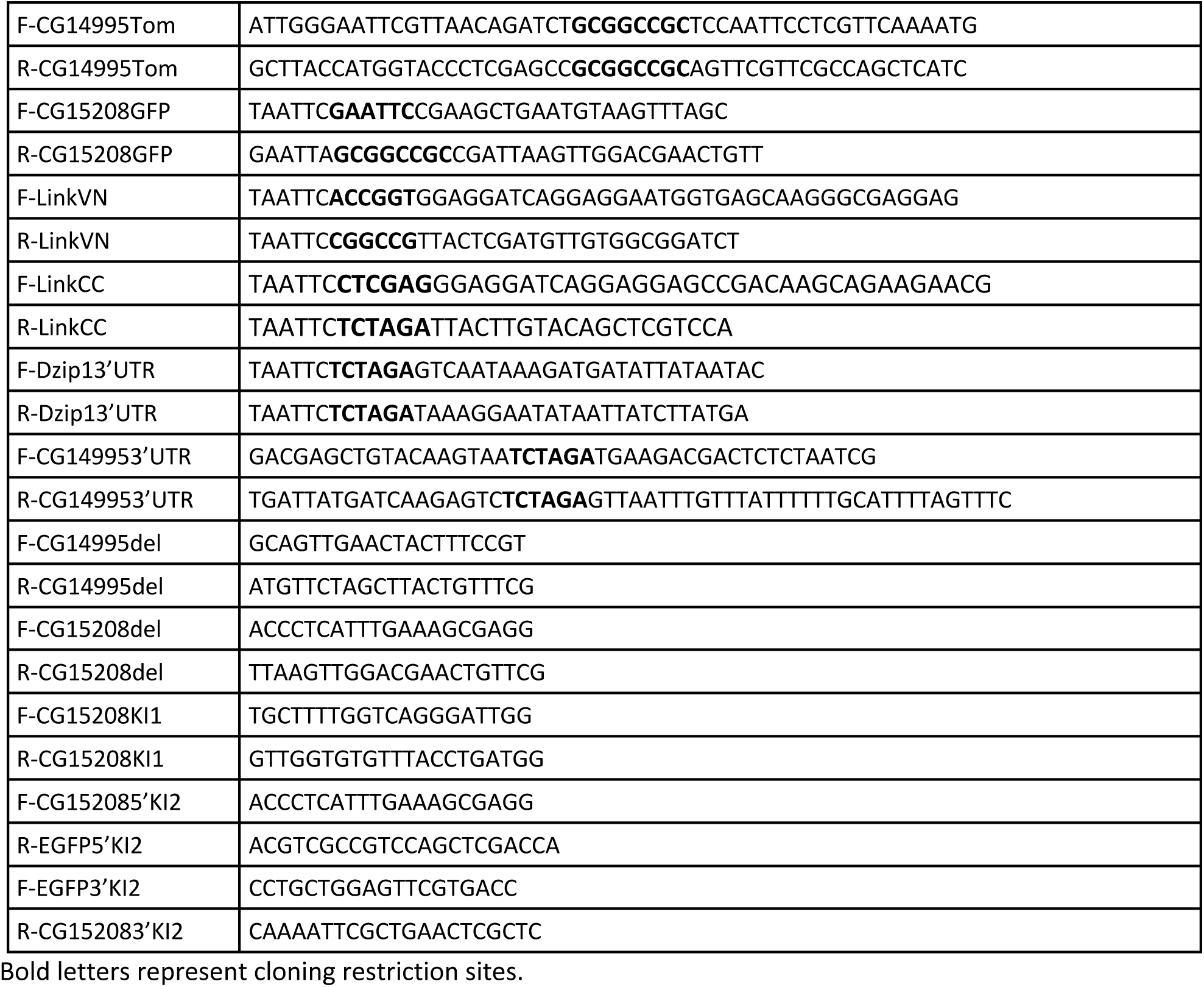
primers used in method section.

